# Deep brain stimulation reduces subthalamic nucleus pathological dynamics and rescues gait deficits associated with dopamine loss

**DOI:** 10.64898/2026.03.17.712325

**Authors:** Leo Steiner, Radu Darie, Audrey Lindsay, Hua-an Tseng, Ingrid van Welie, Xue Han

## Abstract

The Subthalamic Nucleus (STN) regulates movement and is an important clinical target for deep brain stimulation (DBS) in Parkinson’s Disease (PD). However, it remains unclear how dopamine loss and DBS influence STN gait encoding. We performed simultaneous recordings from multiple neurons and intermittent DBS in the STN of healthy and dopamine depleted PD mice during voluntary locomotion. We found that dopamine loss resulted in gait deficits manifested as altered stride length of both hindlimbs and forelimbs, which were rescued by intermittent DBS. Furthermore, dopamine loss exaggerated movement encoding of STN population dynamics, and elevates individual STN spiking during movement and beta-rhythmic firing at rest. Despite an overall increase in the fraction of neuron activated by movement, individual neurons gait encoding properties remain similar between healthy and PD mice. While DBS suppressed firing in both healthy and PD mice, it selectively reduced STN beta-rhythmic spiking, desynchronized STN networks, and rescued gait deficits associated with the loss of dopamine. These results suggest that pathological activation and beta synchronization of the STN contributes to motor deficits related to PD, and DBS-induced reduction of beta rhythmic spiking and STN network desynchronization contribute to the therapeutic effects of DBS in PD.

## Introduction

Parkinson’s Disease (PD) is attributed to the progressive degeneration of midbrain dopaminergic neurons, resulting in cardinal motor symptoms such as akinesia, dyskinesia, tremor, gait disturbance, and postural instability^1–6^. Lower extremity and axial symptoms, specifically gait and postural instability, often progress more rapidly than upper extremity tremors^4,7–9^. Current therapies, including dopamine replacement and deep brain stimulation (DBS), are effective for reducing tremor, bradykinesia, and rigidity, but they have mixed effects on gait and sometimes worsens gait^4,10–19^. Converging evidence suggests a progressive worsening of gait after subthalamic nucleus (STN) DBS, with up to 42% of patients reporting gait deterioration after 6 month of DBS electrode implantation^20^. Gait speed and variability are important aspects of walking, which are extensively regulated by the spinal cord, and broad cortical and subcortical brain areas. While the spinal central pattern generators are critical for regulating gait^21,22^, gait related neural dynamics are broadly detected across the cortical-basal ganglia-thalamic motor circuits, including the motor cortex^23–25^, STN^26–30^, and dorsal striatum^31–33^.

One common target of DBS in PD is the STN^34,35^. The STN receives prominent inhibitory inputs from the globus pallidus externa (GPe)^36,37^ and broad excitatory cortical inputs from associate and limbic areas^38,39^. STN sends excitatory projections to all other nuclei within the basal ganglia^40,41^. These anatomical connections follow the general topographic organization of the basal ganglia, mapping onto the dorsal, medial and ventral subregions of the STN^40,42–44^. STN regulation of movement has been well characterized in humans and nonhuman primates. Around 30-50% of the dorsal STN neurons are activated by passive or active contralateral arm, leg and other joint movement, and do not appear to form well defined anatomical clusters^45–47^. While difficult to target due to its small size in mice, estimated to be ∼0.09 mm^3^ in volume^48,49^, a recent single unit recording study revealed that firings in many STN neurons are modulated by stepping movement in mice^27^. In primates, loss of dopamine in PD mildly increases STN firing rate^50^, disrupts STN arm movement encoding^51^, and reduces STN specificity by increasing response to ipsilateral joint movement^52^. In addition to its role in regulating movement kinematics, STN neurons, being part of the “indirect pathway” of the basal ganglia^37^, are engaged in action selection and movement cancellation^46,53–55^. These findings are in line with the observation that surgical resection of STN, performed historically to treat Parkinsonism, may reduce movement inhibition and promote dyskinesia^50,56,57^. STN is also involved in various nonmotor functions, particularly the medial and ventral subregions^40,42–44^. For example, lesion and stimulation of the ventromedial STN alter limbic functions^58^, including reward processing^59^, proactive inhibition^60^, attention, and arousal^61^.

Synchronized beta-frequency (∼13-30 Hz) STN dynamics have been broadly captured by local field potential (LFP) recordings in human patients and PD animal models^16,19,62,63^, and serve as a clinical PD biomarker^5,64–67^. Under healthy conditions, beta oscillations are thought to serve as a “maintaining status quo” signal^65,68^, which is suppressed during movement initiation and action^69^ and augmented during movement stopping^70,71^. Consistent with a role in moment-to-moment movement regulation, STN LFP beta power was recently shown to be modulated by gait cycles^26,28,30^. Exaggeration of beta oscillations thus has been theorized to inhibit movement, particularly relevant to PD akinesia.

While the therapeutic mechanisms of DBS remain unclear, DBS has been shown to activate local and passing axons and suppress local neurons, both of which could cause functional “informational lesion” to reduce pathological circuit dynamics^72–74^. To understand how DBS and dopamine loss impact individual STN neurons and STN population dynamics related to gait, we designed and fabricated custom multisite micro-electrode probes that allowed for simultaneous recording of tens of STN neurons in the small mouse STN. We systematically characterized individual STN neuron spiking dynamics, STN network synchrony, and the effect of DA loss in voluntarily locomoting mice. In addition, the custom microelectrode probes contained integrated electrical micro-electrode stimulation sites, enabling us to probe the effect of STN DBS on DA-dependent gait pathology and STN gait encoding.

## Results

### Multisite STN recording during voluntary locomotion with precision gait analysis

To characterize the effect of DA loss and DBS on STN dynamics during locomotion, we simultaneously recorded multiple STN neurons using custom linear micro-electrode arrays, while mice were head fixed and voluntarily locomoting on a disk treadmill (**Fig. 1A-D**). Briefly, we first surgically implanted a metal head plate for head fixation and performed a craniotomy over STN (Methods). To investigate the effect of DA loss, about half of the mice additionally received neurotoxin 6-OHDA infusion in the medial forebrain bundle (PD mice, n=9 mice), while the other half did not (Healthy mice, n=11 mice). Upon recovery from the surgery, mice were habituated to the recording setup, and a single acute recording session was performed in each mouse to allow for verification of STN targeting via histology. Specifically, on the day of recording, we first dipped the electrode array in cell-tracker dye that allowed for later visualization of electrode track during histological validation (**Fig. 1E**). We then removed the dura from the previously made craniotomy, and slowly inserted the electrode array until the deepest channel reached STN. In general, STN was about 4.5 mm ventral to the pia, beneath the thin Zona Incerta area, and electrophysiologically identified by neurons tonically firing at ∼10-30 Hz^51,75–78^. Three to five days after the recording session, histology was performed to visualize the electrode tract marked by cell tracker dye. Of the 20 mice recorded, electrode tracts were visible in the STN in 14 mice. Two of these mice showed little movement (Methods), and thus were excluded from further analysis. All subsequent analysis was performed on 12 mice with STN verified both physiologically during recording and histologically post-recording (n=6 healthy mice, and n=6 PD mice).

**Figure 1:**
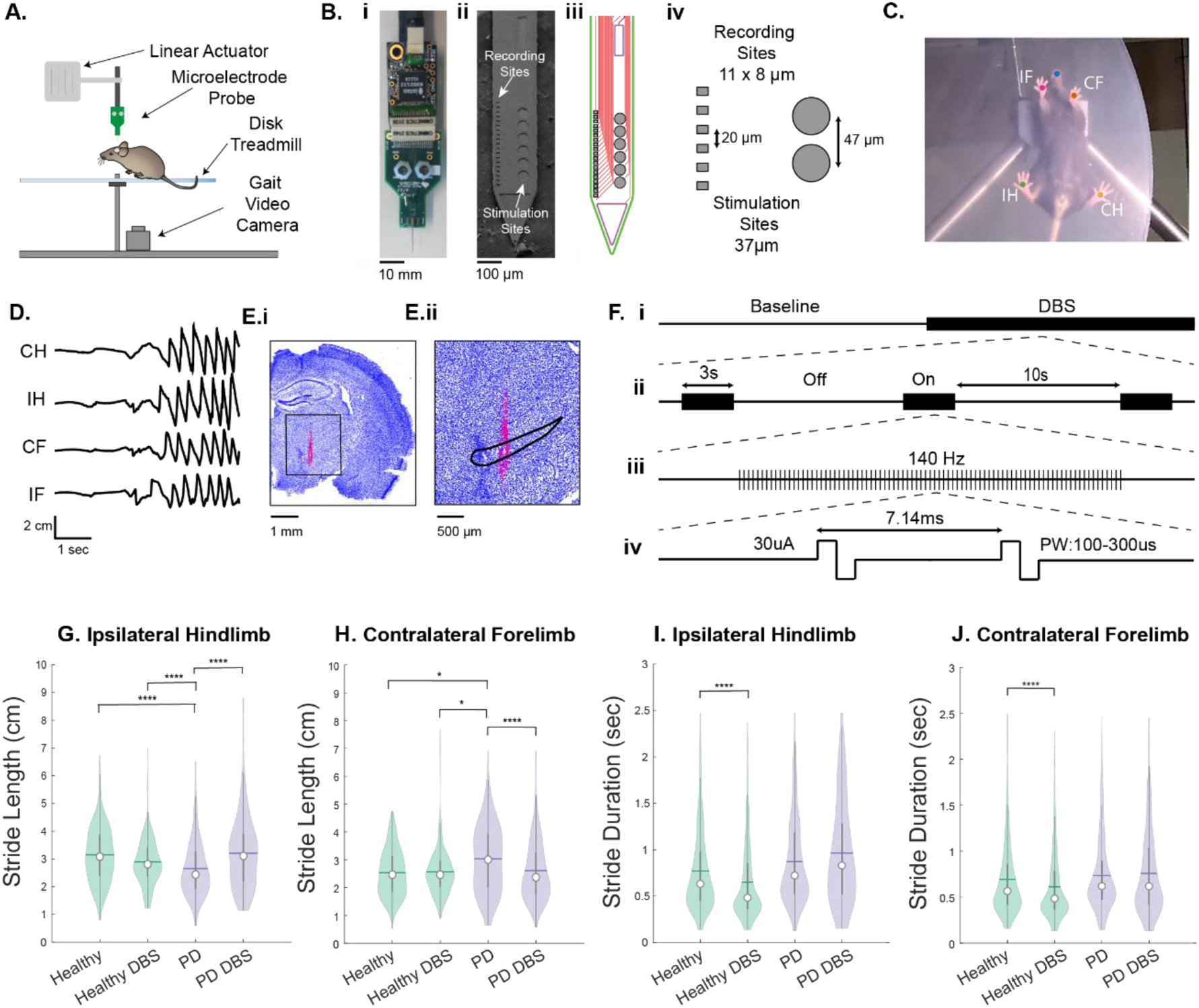
PD mice show asymmetrical stride length of both hindlimbs and forelimbs, which are both rescued by DBS. (**A**) Schematics of experimental setup. (**B**) Design of Linear electrode array, showing (**i**) photograph of the whole probe, (**ii**) electron microscope image of the recording and stimulation sites, (**iii**) PCB wiring, and (**iv**) schematic. (**C**) An example video frame showing individual limbs identified. “IF”, ipsilateral forelimb; “CF”, contralateral forelimb; “IH”, ipsilateral hindlimb; “CH”, contralateral hindlimb. (**D**) Example traces of limb position. (**Ei**) An example histological brain slice stained with DAPI (blue), showing electrode tract (red). (**Eii**) Zoom in of **Ei** with STN outlined (black). (**F**) DBS protocol and pulse parameters. (**G-J**) Population stride length across healthy and PD groups for ipsilateral hindlimb (**G**), contralateral forelimb (**H**) stride lengths, and stride duration for ipsilateral hindlimb (**I**) and contralateral forelimb (**J**). Quantifications in G-J are visualized as violin plots with the outer shape representing the data kernel density.GLME, *, p<0.05. **, p<0.01; ***, p<0.005. ****, p<0.001.

The custom electrode arrays contained 19 recording sites, 20 μm apart, spanning 380 μm in total (**Fig. 1B**). This spans the dorsoventral axis of the mouse STN, which we estimated to be approximately 300 μm in height and ∼0.09 mm^3^ in volume using Allen Brain Institute datasets^49^. Additionally, they had 6 stimulation sites, 47 μm apart, centered next to the recording sites. Spike waveforms recorded from all 19 channels were preprocessed in Kilosort4 to identify individual neurons (Methods), resulting in 13.2 ± 4.3 neurons per session recorded (mean ± standard deviation (SD), n=12 sessions in 12 mice). We assigned the location of the spiking neuron to the recording site with the highest amplitude spike waveforms, which yielded 10.4 ± 4.0 spike containing recording sites per session. To capture each animal’s stepping movement, a video camera was positioned under a transparent disk treadmill^79^ to track individual limb movement at 60 frames/sec. The relative positions of all four limbs in each video frame were then extracted using DeepLabCut^80^ trained neural networks (**Fig. 1C-D**).

During each recording session, we first obtained a baseline recording of STN neurons and stepping movement for 10-20 minutes (11.1 ± 4.4 minutes, mean ± SD, n=12 sessions in 12 mice). We then delivered intermittent DBS through all stimulation sites, using 140 Hz, biphasic, charge balanced pulse trains for a total of about 3 minutes per recording (3.4 ± 0.66 minutes). Each pulse train was 3 seconds long, with 10 second inter-train interval (**Fig. 1F**). DBS pulse width was adjusted empirically in each mouse to be 75% of the minimum pulse width needed to evoke any noticeable movement behaviors (202 ± 104 µs per phase, Methods).

### Unilateral DA loss impaired ipsilateral hindlimb and contralateral forelimb movement, which were rescued by DBS

To assess the effect of unilateral DA loss on voluntary locomotion, we first characterized the rotational behavior in an open field environment at 3-5 days post 6-OHDA injection. PD mice consistently exhibited more ipsilateral rotations than contralateral rotations, and more ipsilateral rotations than healthy mice (**Sup. Fig. 1F**, Kruskal-Wallis, p=0.0158, p=0.024), confirming successful DA depletion as broadly shown previously ^81,82,2^. DA loss was further confirmed via immunohistochemistry against tyrosine hydroxylase, revealing a 62 ± 20% (mean ± SD, n=6 mice) reduction of immunofluorescence in the striatum (**Sup. Fig. 1B**).

Having confirmed the loss of striatal DA in PD mice, we next evaluated how DA loss impacted locomotion and gait under head-fixed experimental conditions. We first identified individual steps of each limb using relative limb positions. Movement bouts were then defined as periods containing 3 or more steps. The onset of a movement bout was identified as the beginning of the first step of any of the four limbs, and offset was the end of the last step. Rest periods were periods without any limb movement. Gait analysis, including stride length and stride duration, was restricted to movement bouts.

All 6 PD mice showed an asymmetry of stride length, with every mouse having shorter ipsilateral hindlimb stride lengths than contralateral hindlimb lengths (**Sup. Fig. 2A**, Kruskal-Wallis, p<0.05 for all mice), and 5/6 PD mice additionally showed a longer contralateral forelimb stride length than the ipsilateral forelimb (**Sup. Fig. 2A**). In contrast, none of the healthy mice showed a difference in the stride length for either the hindlimb or forelimb (**Sup. Fig. 2B**, Kruskal-Wallis, p>0.05 all for mice). Compared to the healthy population, the PD population exhibited a significant reduction in ipsilateral hindlimb stride length and an increase in contralateral forelimb stride lengths **(Fig. 1G,H**, GLME, p=2.1e-5 and p=0.048 respectively). We then assessed asymmetry by comparing the stride length ipsilateral to the recording site to the corresponding contralateral ones. Across PD mice, there was a significant asymmetry for both hindlimbs and forelimbs, whereas healthy mice showed no asymmetry (**Sup. Fig. 2C-F**, Mann-Whitney U, PD hindlimbs p=1.1e-21, forelimbs p=1.1e-16, healthy hindlimbs p=0.101, forelimbs p=0.814).

**Figure 2:**
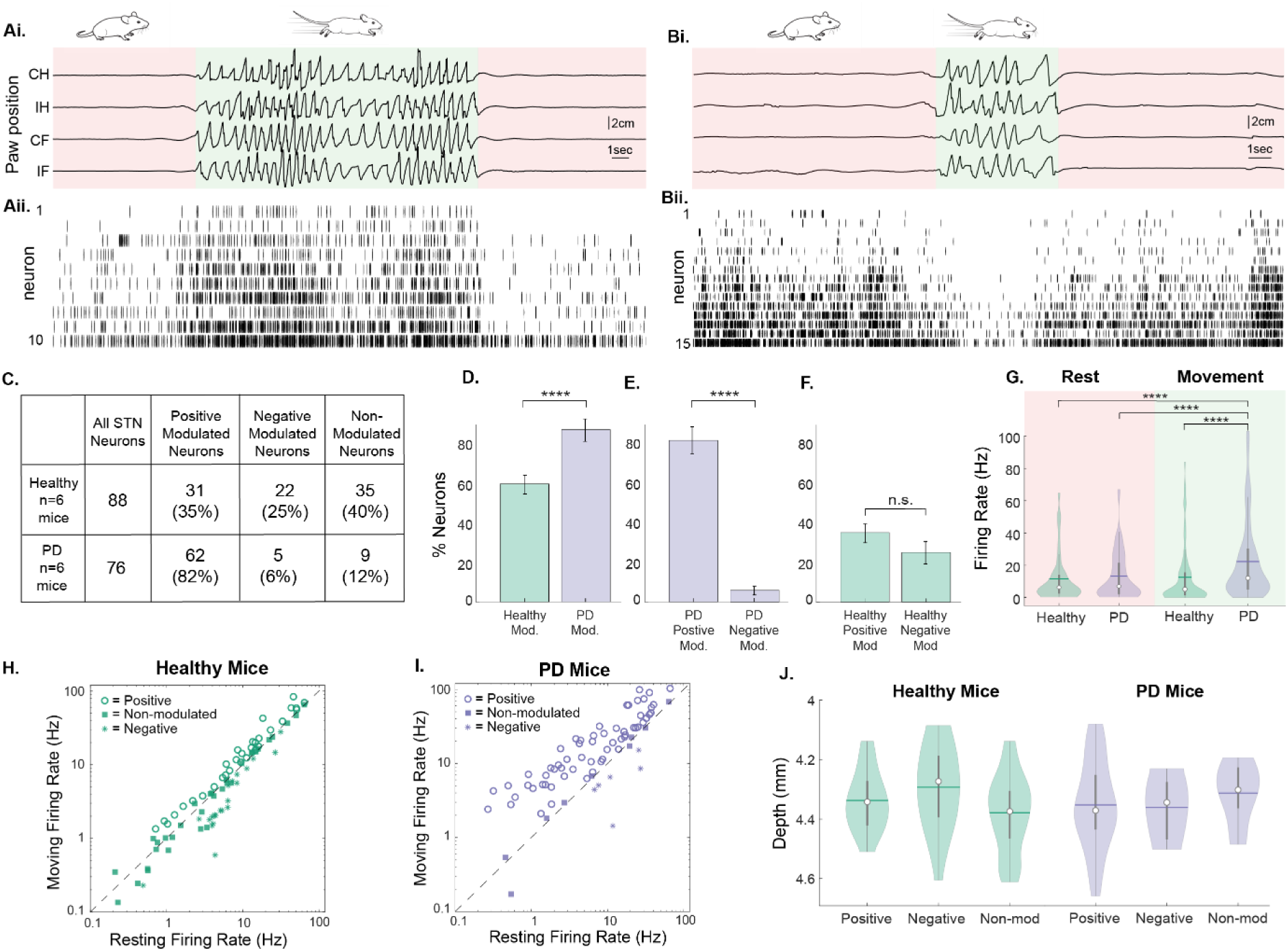
STN neuron activation during movement was exaggerated in PD mice. (**A**) Example limb positions and corresponding spiking from 10 simultaneously recorded STN neurons in a healthy mouse. (**i**) relative limb positions of the four limbs. Green shading indicates movement bouts, and red shading indicates rest. (**ii**) Spike rasters. Neurons were sorted by firing rate. (**B**) Similar to **A**, but for a PD mouse showing 15 simultaneously recorded STN neurons. (**C**) Summary of movement response of neurons in healthy and PD mice. (**D**) The percentage of modulated neurons in healthy and PD groups. Bars represent observed percentage of modulated neurons and error bars are estimated standard error. (**E, F**) The percentage of positively and negatively modulated neurons in PD mice (**E**) and in healthy mice (**F**). (**G**) Firing rate across all neurons at rest and during movement in healthy and PD groups. (**H**) The relationships of firing rates of individual neuron in healthy mice during movement versus at rest. Neurons are labelled by their movement modulation. (**I**) Similar to **H**, but for PD mice. (**J**) Location of recorded neurons in healthy and PD mice, grouped by their movement response. Quantifications in **G** and **J** are visualized as violin plots with the outer shape representing the data kernel density Chi Square *, p<0.05, **, p<0.01. ***, p<0.005. ****, p<0.001.

We next explored how DBS modulates stride length. In 3 of the 6 PD mice, DBS increased ipsilateral hindlimb stride length and reduced contralateral forelimb stride length compared to those without DBS (**Fig. 1F,G, Sup. Fig. 2A**, Kruskal-Wallis, p>0.05), and rescued DA loss-induced asymmetry of both the hindlimbs and forelimbs (**Sup. Fig. 2H, J**, Kruskal-Wallis, hindlimb: p=0.955, forelimb: p=0.993). Interestingly, DBS over compensated hindlimb asymmetry, and rescued forelimb asymmetry across PD population (**Sup. Fig. 2H**). Across the PD population, DBS significantly increased ipsilateral stride lengths and reduced contralateral stride length (**Fig. 1G, H**, GLME, hindlimb: p=4.2e-8, forelimb: p=1.8e-4), and rescued stride length to that observed in healthy mice (**Fig. 1G, H**, GLME, hindlimb: p=0.664, forelimb: p=0.445). In contrast, DBS did not alter stride length of any limb in any individual healthy mice (**Sup. Fig. 2B**, Kruskal-Wallis, p>0.05), or across the healthy population (**Fig. 1G, H**, GLME, hindlimb: p=0.351, forelimb: p=0.126). In addition, most PD mice had significantly longer stride duration than healthy mice, but failed to reach significance on a population level (**Fig. 1I, J**, GLME, p=0.181, p=0.389). However, DBS decreased stride duration in healthy mice (**Fig. 1I, J**, GLME, p=8.6e-4, p=0.003), but not PD mice, suggesting that DBS selectively increased movement speed of healthy mice. Finally, other movement parameters, including bout duration, number of strides per bout, percent of time moving, and the number of bouts per minute were similar between healthy, health DBS, PD, and PD DBS groups (**Sup. Fig. 2S-V**, Kruskal-Wallis, p>0.5).

Together, these results demonstrate that unilateral DA loss impacts gait in head-fixed conditions, consistently reducing ipsilateral hindlimb stride length and increasing contralateral forelimb stride length, leading to movement asymmetry. The impact of DA loss on stride length is consistent with the ipsilateral rotational behavior of the PD animals in the open field. DBS in the STN selectively rescued DA loss mediated changes in stride length and movement asymmetry in PD mice, while increasing overall movement speed in healthy mice without altering other gait parameters.

### DA loss exaggerates STN movement encoding and elevates STN spiking during movement

In addition to the extensive studies of STN dynamics during action selection in non-human primates^54,55,57,70^, recent evidence suggests that STN neurons exhibit heterogeneous dynamics during voluntary movement in mice^27^. We noted that many STN neurons in heathy mice were modulated by movement, either increasing or decreasing firing during movement (**Fig. 2A, B**). To characterize the movement encoding properties of individual neurons, we compared the firing rate at rest versus during movement using a boot strapping method (Methods). We found that 88% of neurons in the PD group were movement modulated, significantly more than the 60% observed in the healthy group (**Fig. 2C, D**, Chi-Square, p=2.3e-9). We then further classified the significantly modulated neurons as either positively modulated, if a neuron fired more during movement than rest, or negatively modulated if it fired less during movement. Interestingly, positively modulated neurons were more prevalent than negatively modulated neurons in PD mice (82% vs 6%, **Fig. 2E**, Chi Square p=3.3e-12). As a result, across all neurons in PD mice, the population firing rate was significantly higher during movement than rest (Wilcoxon signed-rank, p=6.5e-10, n=88 healthy, n=76 PD **Fig. 2G**). In contrast, a balanced proportion of positively and negatively modulated neurons were detected in healthy mice (**Fig. 2F**, 35% versus 25%, Chi Square, p=0.055, n=31 positively modulate, n=22 negatively modulated), and the average population firing rate was similar between movement and rest (Wilcoxon signed-rank, p=0.396, **Fig. 2G**).

It is well recognized that the dorsal subregion of the primate STN is associated with sensorimotor function, whereas the ventral subregions are more linked to associative and limbic function^40,83^. As our electrode arrays span most of the STN along the dorsoventral axis, we examined the relative location of movement modulated neurons along the electrode tracts. We did not observe any spatial clustering of modulated neurons along the dorsoventral axis, either positively or negatively modulated ones (**Fig. 3J**), consistent with some previous findings in non-human primates^84^, but could also reflect the small size of the mouse STN. Together, these results demonstrate that STN neurons exhibit balanced modulation during movement under healthy conditions, and DA loss exaggerate the positive responding during movement, consistent with the theory of pathologically activated indirect pathways in PD^37^.

**Figure 3:**
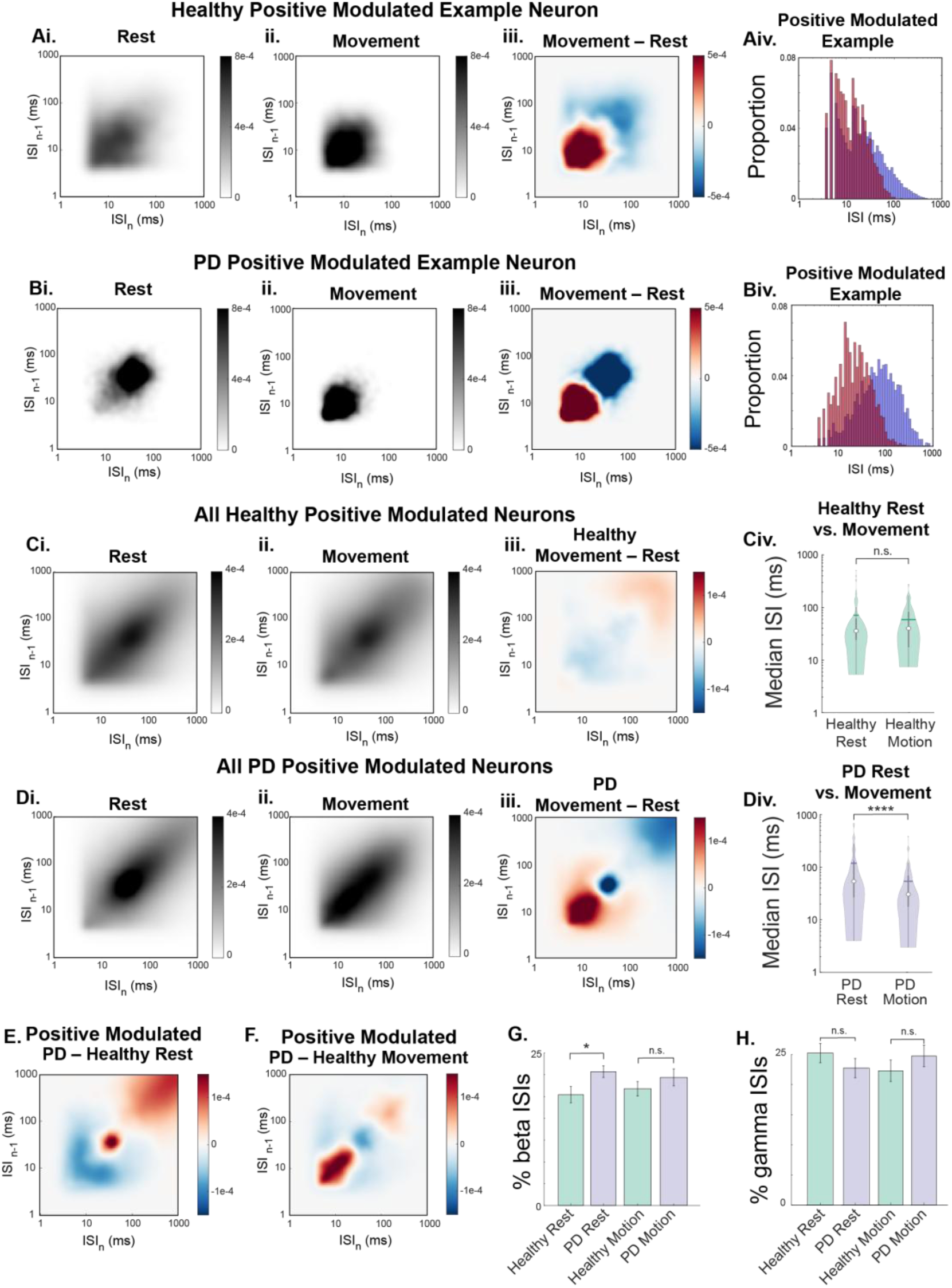
Positively modulated STN neurons in PD mice exhibited an enhanced interspike interval (ISI) specifically at beta frequency at rest. (**A**) ISI return map of an example positively modulated neuron from a healthy mouse at rest (**Ai**), during movement (**Aii**), and their difference (**Aiii**, shown as **Aii** minus **Ai**). (**Aiv**). Histogram of the ISI distribution of the neuron during rest (blue) versus movement (red). (**B**) Similar to **A**, but for an example positively modulated neuron from a PD mouse. (**C**) ISI return map of all positive modulated neurons of healthy mice at rest (**Ci**), during movement (**Cii**), and their difference (**Ciii**, shown as **Cii** minus **Ci**). Median ISI from each positive modulated neuron at rest and during movement (**Civ**). (**D**) Similar to **C**, but for all positvely modulated neurons from PD mice. (**E-F**) Difference in the ISI return map between healthy and PD mouse (PD minus healthy) at rest (**E**) and during movement (**F**). (**G**) Percentage of beta ISIs in healthy and PD mice at rest and during movement. (**H**) Similar to **G**, but for gamma ISIs. Quantifications in Civ and Div are visualized as violin plots with the outer shape representing the data kernel density. Bars in G-H represent the mean percentage of beta or gamma ISIs per neuron, and error bars represent standard error. Wilcoxon rank-sum *, p<0.05.

### DA loss selectively increases STN beta-rhythmic spiking at rest

Firing patterns of STN neurons are known to be diverse and sensitive to DA loss ^27,76,77,85,86^. Because 82% (62/76) of neurons were positively modulated by movement in PD mice, we first investigated how DA loss affected their firing patterns. We computed the inter-spike-interval (ISI) distribution for each neuron at rest and during movement. Most individual neurons in healthy mice exhibit a broad range of ISI distribution patterns, with no obvious difference between rest and run (**Fig. 3A**). However, many neurons in PD mice exhibited a prominent peak around beta frequencies (33-76 ms, 13-30 Hz) at rest, and a peak around gamma frequencies (10-33 ms, 30-100 Hz) during movement (**Fig. 3B**). Higher frequency firing in PD mice during movement is consistent with most neurons being positively modulated.

As movement influences spiking patterns, we first examined the ISI distributions across positively modulated neurons in health and PD mice. In healthy mice, ISI distribution remained largely identical at rest versus during movement (**Fig. 3C**, Kolmogorov-Smirnov (KS) Test of ISI cumulative distribution function, p=0.9655). In contrast, in PD mice, the ISIs drastically shifted from beta to gamma frequency range (**Fig. 3D**, KS test p=0.011). Further comparison between healthy and PD mice revealed a prominent difference in ISI peak distribution at beta frequencies at rest (**Fig. 3E**,), and at gamma frequencies during movement (**Fig. 3F**). To quantify the difference in beta versus gamma rhythmic firing, we computed the fraction of ISIs within the beta or gamma range for each neuron, and compared across all neurons in the two cohorts. We found that beta-frequency ISIs in positively modulated neurons in PD mice were elevated at rest compared with healthy mice, but not during movement (**Fig. 3G**). During movement, there was no significant change at either beta-frequency nor gamma-frequency ISIs (**Fig. 3H**).

While a majority of neurons were positively modulated in PD mice, healthy mice showed a balanced proportion of movement modulation, including many that were negatively modulated. Further evaluation of the ISI patterns in negatively and non-modulated neurons in healthy mice revealed similarly diverse ISI patterns as in positively modulated ones during movement versus rest (**Sup. Fig. 3A, C**). Because of the balanced proportion of positively and negatively modulated neurons in healthy mice, across all neurons, there was little change in ISI distributions between rest and movement (**Sup. Fig. 3E**). Because majority of the neurons were positively movement modulated, across all neurons in PD mice, ISI distribution followed that of the positively modulated ones, exhibiting significantly greater beta-rhythmic firing at rest than healthy mice (**Sup. Fig. 3J**, Wilcoxon rank-sum p= 0.020). Furthermore, consistent with a higher proportion of positive modulated neurons in PD mice than healthy mice, gamma-rhythmic firing during movement across all neurons in PD mice was more prominent than in healthy mice (**Sup. Fig. 3L**, Wilcoxon rank-sum p = 0.027). Overall, these results demonstrate that DA loss selectively exaggerated individual STN neuron’s beta-rhythmic firing at rest, but not during movement. An increase in the fraction of STN neurons showing positive response to movement after DA loss leads to exaggerated gamma-rhythmicity firing of the STN population dynamics firing during movement.

### STN neurons encode gait, and DA loss produces minimal effect on individual STN neuron gait encoding

STN LFP beta power was recently shown to fluctuate with gait cycle in PD patients^26^ and individual STN neurons encode gait in mice^27^. We found that spiking in many neurons in both healthy and PD mice was modulated by the gait cycle (**Fig. 4A-B)**. To determine how DA loss influences STN neuron gait encoding, we computed the relative timing of spikes to the phase of each limb gait cycle (**Fig. 4C-F**). In many neurons, in both healthy and PD mice, we noted that spiking occurred preferentially during the stance phase (**Fig. 4Civ-Fiv**). To determine whether a neuron encodes gait, we examined the relative timing of spikes to gait phase (Spike-gait phase locking, Methods, **Fig. 4G**). We found that most neurons in both cohorts (76% in healthy and 93% in PD mice) exhibited spike-gait phase locking to at least one limb, with more spikes during the stance phase (**Fig. 4H**). In particular, more neurons in PD mice encoded gait of the impacted contralateral forelimbs, but not the other three limbs, compared to those in healthy mice (**Fig. 4H**, Chi Square p=8.3e-4 for contralateral forelimb, and p>0.05 for all other limbs).

**Figure 4:**
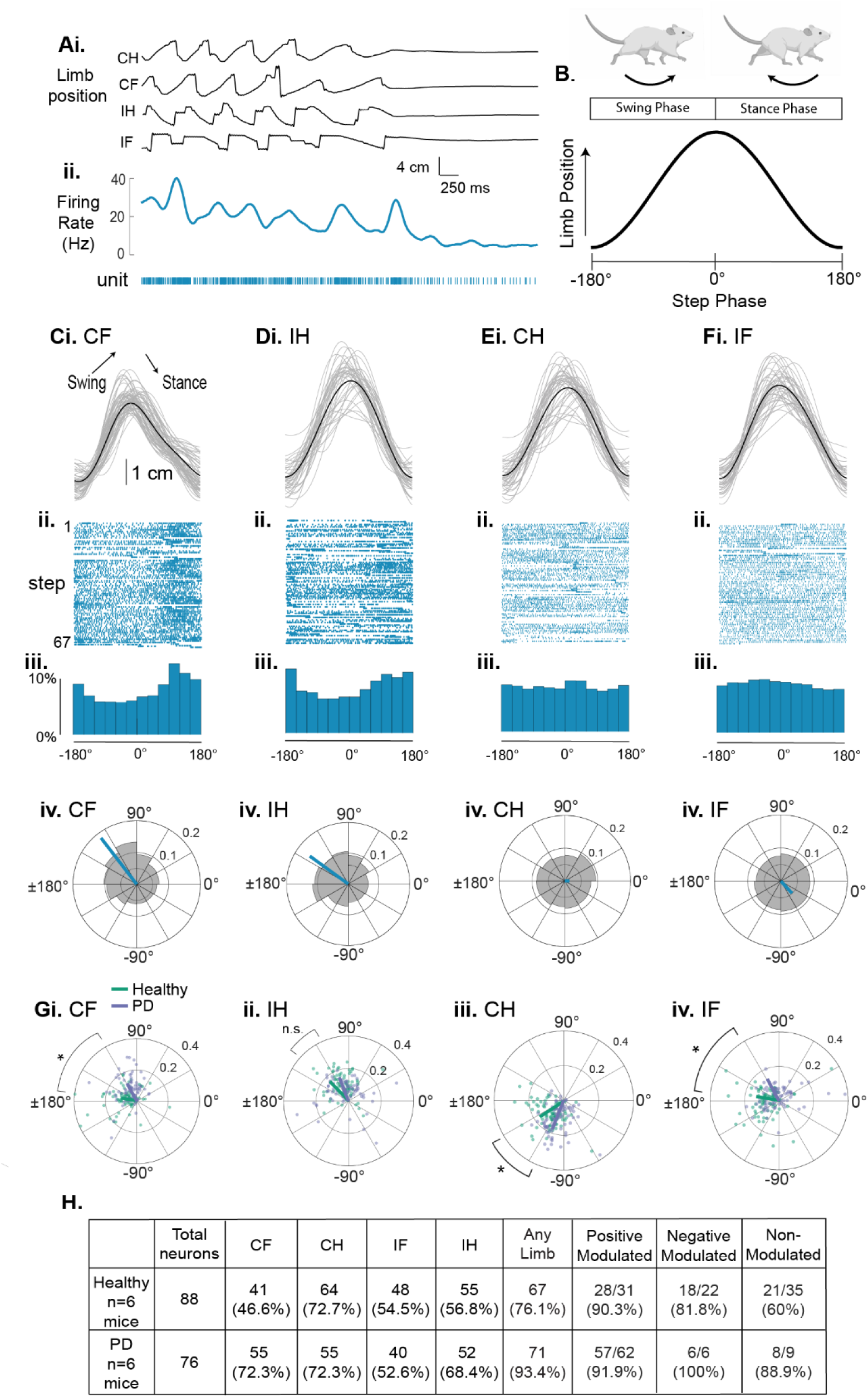
Neurons in PD mice show similar gait cycle dependent firing as in healthy mice. (**Ai**) Example limb traces at the end of a movement bout. CH: contralateral hindlimb, IH: ipsilateral hindlimb. CF: contralateral forelimb, IF: ipsilateral forelimb. (**Ai**) The firing rate and spike timing of an example neuron recorded during **Ai**. (**B**) Illustration of the swing phase (-180° – 0°) and the stance phase (0° – 180°) of a gait cycle. The peak of the gait cycle is defined as 0°. (**C**) Spike raster of an example neuron in healthy mice, rescaled to the start and end of CF gait cycles. (**Ci**) gray: individual gait cycles. Black: averaged gait trace. (**Cii**), corresponding spiking from an example neuron. Each row represents spiking during a gait cycle. (**Diii**), Average spike rate across all gait cycles. (**Div**) Spike-gait phase relationships of the neuron in **C** (Shade area represents the spike phase distribution, and the blue line indicates the mean phase vector). (**D-F**) The same neuron as in **C**, but aligned to the gait cycle of the ipsilateral hindlimb (**D**), contralateral forelimb (**E**), and ipsilateral forelimb (**F**). (**G**) Spike-gait phase relationships of all neurons for contralateral forelimb (**Gi**), ipsilateral hindlimb (**Gii**), contralateral hindlimb (**Giii**), and ipsilateral forelimb (**Giv**) (Each dot represents the averaged phase vector length and phase angle of an individual neuron from healthy (green) or PD (purple) mice). (**H**) Spike-gait phase relationship across all neurons. *, p<0.05..Rayleigh Statistics comparing the populations. Polar comparison of vector power and direction in **G**, Watson Williams test, p<0.05.

To further characterize spike-gait phase locking, we computed the preferred phase and phase-locking strength for each neuron and then compared across all neurons in PD versus healthy mice. We found that while the strength of phase locking was similar between healthy and PD neurons, the preferred phase showed significant but negligible shift towards the stance phase in 3 limbs (**Fig. 4G**, nonparametric Watson-Williams, CF p=2.2e-6, IH p=0.92, CH p=0.006, IF p=0.01). Further comparison of the positive, negative, and non-modulated neuron subgroups between PD and healthy mice revealed no difference in phase locking strength or in preferred phase (**Fig. 4H**). In summary, most neurons in healthy and PD mice fire preferentially during the stance phase of the gait cycle, and DA loss exaggerated the fraction of neurons encoding the contralateral forelimb movement.

### DBS reduces STN firing rate in both healthy and PD mice regardless of movement state, but selectively reduces PD mice beta-rhythmic firing at rest without altering gamma rhythmic firing during movement

While electrical stimulation interferes with real time recordings, we reliably recorded spiking around 5 ms after the stimulation pulse offset by minimizing the electrical stimulation artifact (**Methods, Fig. 5A-B**). Thus, during intermittent DBS, we examined spike rates during the 10 s intervals between stimulation pulse trains, excluding the 5 ms immediately after each pulse train offset, resulting in approximately 3 minutes of DBS periods analyzed (3.4 ± 0.66 mins). During intermittent DBS, each individual neuron’s spike waveform shapes remained the same as that during the 10-20 minutes baseline period immediately before DBS onset (**Fig. 5C-D**, Cosine Similarity>0.90 for all neurons in all 12 mice). Comparing to the baseline, DBS significantly reduced firing rates during both rest and movement, and in both healthy and PD mice (**Fig. 5E-F**, Wilcoxon signed-rank, at rest, healthy p=1.93-4, PD p=0.029. During movement, healthy p=6.4e-6, PD p= 0.007).

**Figure 5:**
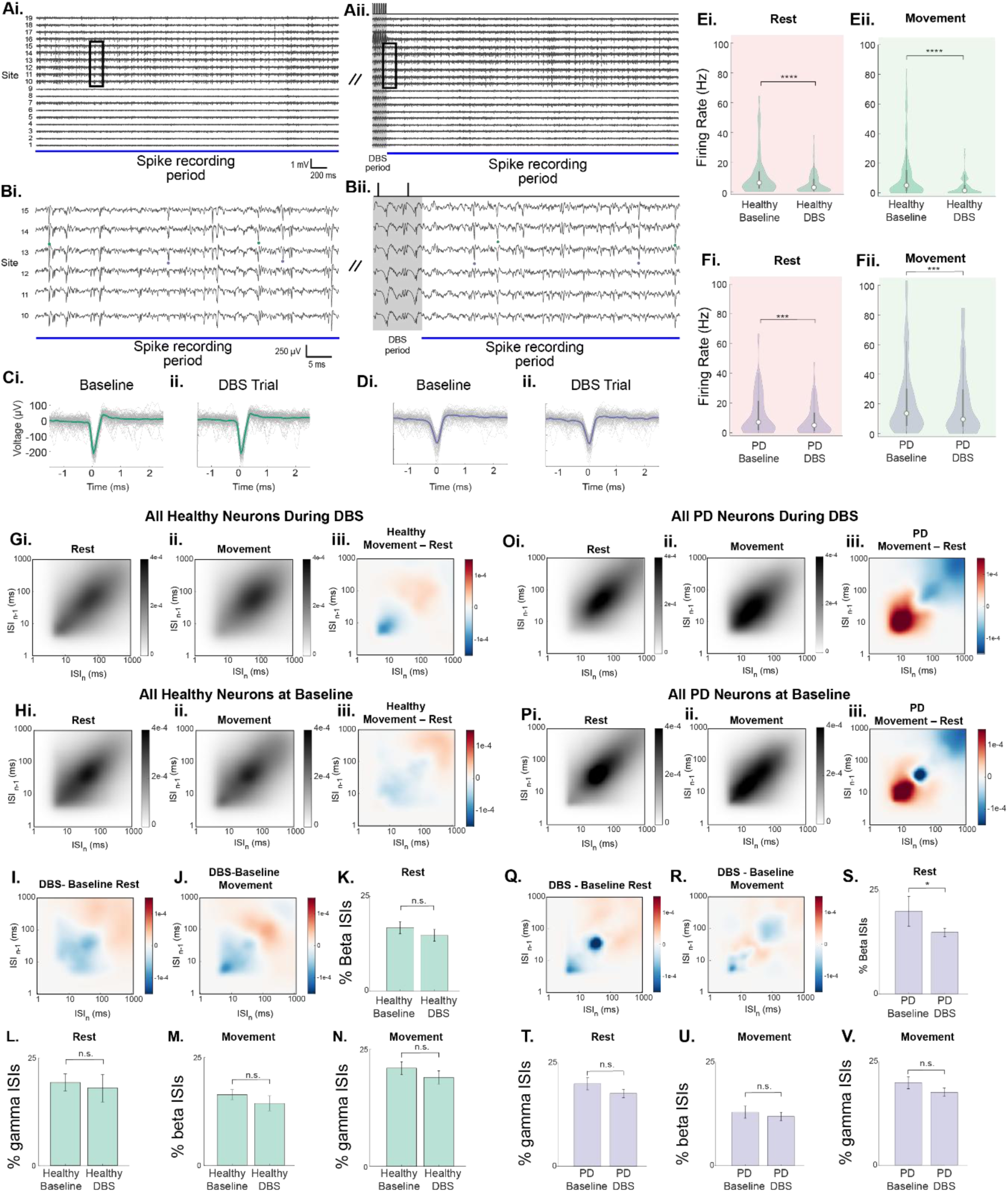
DBS reduces beta-rhythmic ISIs at rest in PD mice. (**A**) An example recording showing 19 sites during baseline period (**Ai**) and at the end of DBS period (**Aii**). (**B**) Zoomed in views of periods highlighted by black boxes shown in **A**. All traces shown are high-pass filtered at 300Hz. Shaded region in **Aii**, **Bii**, indicates periods excluded from analysis, due to artifacts from DBS. Dots indicate spikes from two neurons (green and blue, marked on sites 14 and 13 respectively) during the baseline period and DBS. (**C**) Spike waveforms from green-labeled neuron in **B**, during baseline period and during DBS trial (gray: individual waveform, n=100 randomly chosen waveforms; green: averaged waveform) (**D**) Similar to **C**, but for blue-labeled neuron in **B**. (**E**) Firing rate of STN neurons in healthy mice during baseline and DBS, when mice were at rest (**Ei**) or during movement (**Eii**). (**F**) Similar to **E**, but for STN neurons in PD mice. (**G**) Population ISI return map from neurons in healthy mice during DBS, when the mice were at rest (**Gi**), during movement (**Gii**), and their difference (**Giii,** shown as **Gii** minus **Gi**. (**H**) Similar to **G**, but during baseline period. (**I-J**) The difference in ISI return map between DBS and baseline period (DBS minus baseline) when the mice were at rest (**I**) or during movement (**J**). (**K**) Percentage of ISIs at beta frequency in healthy mice at rest during baseline and DBS. (**L**) Similar to **K**, but for ISIs at gamma frequency. (**M-N**) Similar to **K** and **L**, but for healthy mice during movement. (**O-V**) Similar to **G-N**, but for neurons in PD mice. Bar plots represent the mean percentage of beta or gamma ISIs per neuron, and error bars represent standard error. **K-N** and **S-V**, *, Wilcoxon Signed-Rank p<0.05, Healthy N=88 neurons, PD N=76 neurons.

Further evaluation of the firing patterns across all neurons revealed that DBS did not alter the ISI distribution of healthy mice either at rest or during movement (**Fig. 5G-J**), and did not alter the fraction of beta or gamma frequency ISIs during either movement state (**Fig. 5K-N**). In contrast, DBS in PD mice reduced beta-frequency ISIs at rest (**Fig. 5Q, S**, Wilcoxon signed-rank p=0.029), but not during movement (**Fig. 5U**, p=0.099). Furthermore, DBS in PD mice did not alter gamma-frequency ISIs either at rest or during movement (**Fig. 5T-V**). Similar changes were also detected when comparing only positively modulated neurons (**Sup. Fig. 4A-B**). Finally, beta-frequency ISIs were reduced during movement in PD mice, but not healthy mice (**Sup. Fig. 4C**), consistent with the observation of reduced LFP beta rhythmicity in patient STN during repeated movements^87^.

In summary, while DBS reduced overall firing rates in both healthy and PD mice regardless of movement state, DBS selectively reduced beta-rhythmic firing in PD mice at rest. As DA loss only enhanced beta-rhythmic firing at rest but not during movement, without affecting gamma-rhythmic firing either during rest or movement (**Fig. 3G-H**), these results demonstrate a specific effect of DBS on rescuing DA-sensitive pathophysiology of individual STN neuron via modulating the beta-rhythmic firing at rest. Furthermore, these results are consistent with the general observation of DBS in reducing STN population beta dynamics related to bradykinesia and rigidity^87^.

### DBS selectively reduces STN network synchrony in PD mice at rest, but not during movement

Increased STN synchrony, measured as augmented LFP oscillation power, has been widely associated with motor deficits in PD^88,89^. As LFP signals are dominated by synaptic activity^90^, it remains unclear whether DA loss leads to synchronized STN spiking output. With the ability to record multiple STN neurons, we examined spike synchrony between simultaneously recorded spike trains by computing pairwise cross-correlation (Methods). As movement influences STN dynamics, we first computed the correlation coefficients at rest. Before DBS, the correlation coefficients between most neuron pairs were the highest at zero lag (Pearson’s Correlation) in both healthy (**Fig. 6A**) and PD mice (**Fig. 6D**), consistent with STN neurons being driven predominantly by shared synaptic inputs^91^. However, during DBS, while most neuron pairs in healthy mice exhibited a similar temporal correlation profile as that before DBS (**Fig. 6B**), many pairs in PD mice exhibited drastically reduced correlation (**Fig. 6E**). To estimate network synchrony, we computed the mean Pearson’s correlation across all simultaneously recorded neuron pairs of a session (**Fig. 6C, F**). At rest, DBS did not alter STN network synchrony in healthy mice (**Fig. 6Civ**, Wilcoxon signed-rank, p=0.437), but significantly reduced synchrony in PD mice (**Fig. 6Civ**, Wilcoxon signed-rank, p=0.0312).

**Figure 6:**
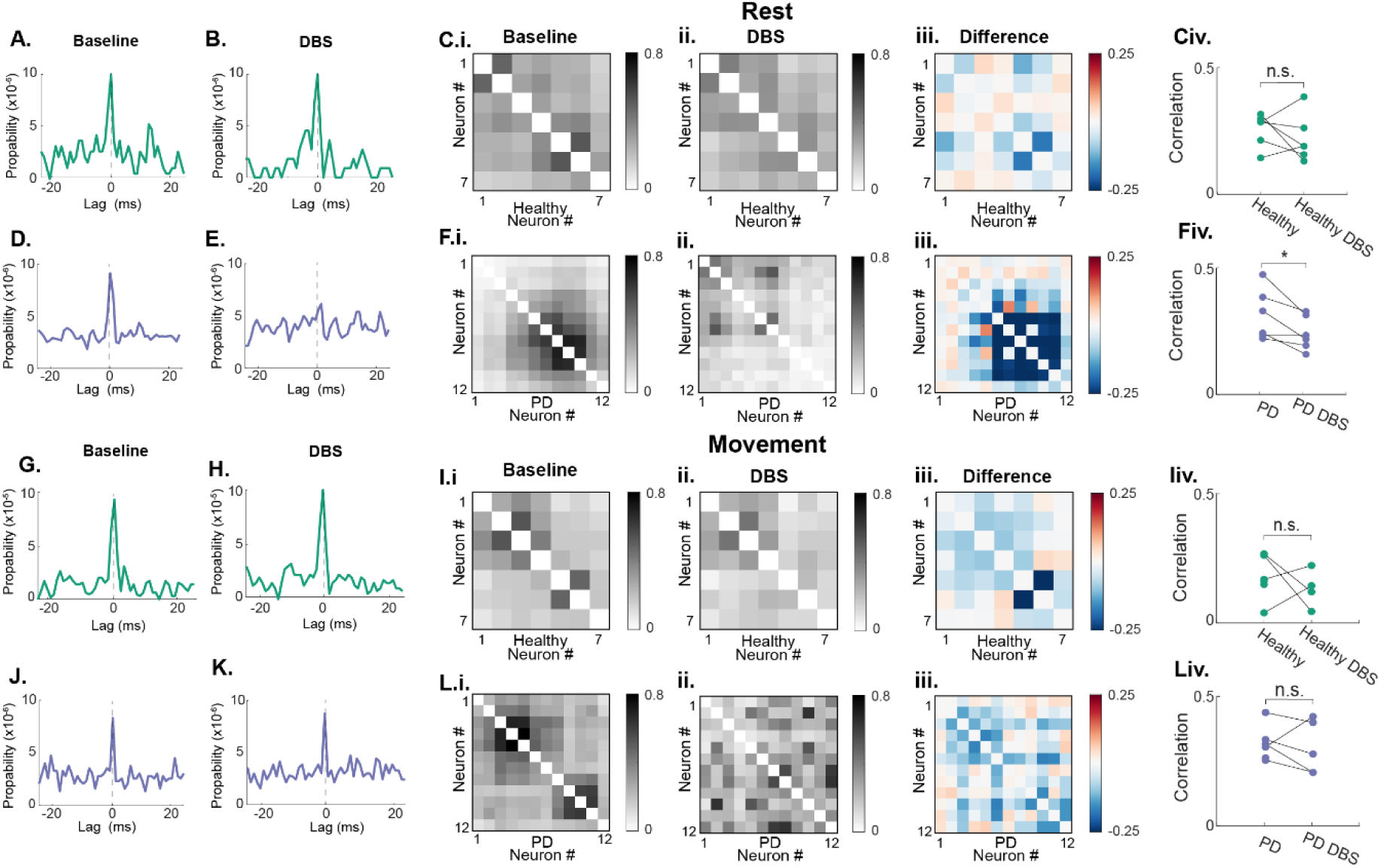
Dopamine loss in PD mice increases STN spike synchrony at rest, and DBS reduces STN synchrony in PD mice at rest. (**A**) Coefficients of cross-correlation between an example pair of neurons from healthy mice at rest, during baseline before DBS. (**B**) Similar to **A**, but for the same pair during DBS. (**C**) Correlation coefficients of all pairs of neurons from an example recording session in a healthy mouse at rest during baseline (**Ci**), DBS (**Cii**), and their difference (**Ciii**, shown as **Cii-Ci**) (**Civ**) Mean correlation coefficients across neurons in healthy mice during baseline versus DBS. (**D-F**) Similar to **A-C**, but for PD mice. (**G-L**) Similar to **A-F**, but during movement period. Wilcoxon signed-rank *, p<0.05.

During movement, DBS did not alter STN synchrony in either healthy or PD mice (**Fig. 6G-L**, p=0.875, p=0.625), underscoring that DBS effects were specific to the resting period. Interestingly, synchrony was significantly higher in PD mice than healthy mice during movement before DBS, but not at rest (**Sup. Fig. 5C**, GLME p= 7.1e-4, p=0.252), consistent with the general observation of augmented synchrony in PD^90^. As DBS effect is restricted to resting period, the elevated network synchrony observed during movement in PD mice appears to be resistant to DBS-mediated modulation, further highlighting a state-dependent effect of DBS. As rest periods accounted for the majority of the recording time, DBS reduced STN synchrony across the entire recordings in PD mice (regardless of movement), but not in healthy mice (**Sup. Fig. 5Eiv**, Wilcoxon signed-rank, p=0.0312). To rule out synchrony changes due to firing rate, additional analysis using Spike Time Tiling Coefficient (STTC) as a measure of network synchrony revealed results consistent with Pearson’s correlation (**Sup. Fig. 5H-K**). Thus, DBS selectively reduces STN synchrony in PD mice at rest, without altering synchrony during movement in PD mice, or during either movement state in healthy mice. These results provide direct experimental evidence that DBS selectively reduce STN synchrony that is relevant for DA-loss mediated circuit pathology.

## Discussion

Through recording many STN neurons simultaneously, we characterized the effect of intermittent DBS and DA loss on the dynamics of individual neurons and the STN network in mice during voluntary locomotion. We detected a nearly balanced proportion of STN neurons that were activated versus suppressed during locomotion in healthy mice. In contrast, in PD mice, STN neurons were predominately activated during movement, and showed exaggerated beta-rhythmic firing at rest. However, the gait encoding features of individual STN neurons remained similar between healthy and PD mice. Intermittent DBS rescued DA loss mediated gait deficits and restored gait symmetry in PD mice. Consistent with the observed gait improvement, DBS selectively reduced beta-rhythmic spiking at rest and desynchronized STN network in PD mice, even though it non-selectively reduced firing in both healthy and PD mice. The observed effects of DBS, including reduced pathological activation of STN, decreased beta rhythmic firing at rest, and desynchronization of the STN network, support beta reduction and network desynchronization as DBS therapeutic mechanisms.

### Unilateral DA loss leads to shorter stride length and asymmetric gait, which are rescued by DBS

One key clinical symptom of shuffling of gait is shorter stride length^92^. Through precision gait analysis, we observed the same phenomenon in the unilateral 6-OHDA lesioned hemi-Parkinsonian mice, consistent with previous observation of speed-dependent stride length reduction in this model^93^. Specifically, we observed stride length reduction of the ipsilateral hindlimb under the head-fixed condition and increased ipsilateral rotation in open field. We also detected STN hyperactivity during movement in PD mice, which could lead to exaggerated activation of the ipsilateral SNr and downstream brainstem-spinal motor circuits that have recently been implicated in driving ipsilateral rotation via reducing the range of ipsilateral limb motion and stride length^81,94^. Furthermore, we also found that DBS suppresses STN hyperactivity, suggesting that DBS restoration of impaired stride length may be through inhibiting STN activation of the SNr. Interestingly, we found an increase in the contralateral forelimb stride length in head-fixed PD mice (**Fig. 1**), rather than reduced ipsilateral forelimb length as observed by Cregg et. al^81^. This likely reflects a compensatory behavior under the head-fixation condition. Thus, our results highlight the potential of using the 6-OHDA lesioned mouse model for mechanistic studies of PD gait.

### DA loss exaggerates STN activation during movement

We found a higher proportion of positively modulated neurons during movement in PD mice than healthy mice, which is most likely related to the reduced inhibition from the inhibitory GPe that is known to be hypoactive in PD^95^. GPe neurons synapse on the proximal dendrites of STN neurons and powerfully regulate STN spiking. A reduction in GPe inhibition could thus increase the somatic depolarization effect of the cortical and thalamic excitatory inputs that are typically on distal dendrites^96–98^, and thereby increase STN activation by cortical and thalamic sensorimotor signals.

### DA loss specifically increases individual neuron’s beta-rhythmic firing at rest and DBS reduces this pathological beta

Beta oscillations across cortical-basal ganglia-thalamic motor circuits is an established biomarker for PD^65,67,99,100^, though the exact relationships between pathological beta of each brain circuit and PD symptoms remain unclear. We quantified the relative change in STN firing at beta versus gamma frequencies, during movement versus resting, in PD versus healthy mice, with and without DBS. We detected a specific increase in beta-rhythmic firing at rest in PD mice, which was reduced by DBS in PD mice only. This selective increase in beta at rest in PD is consistent with the general role of beta oscillations in maintaining status quo, and the exaggeration of such beta in PD could contribute to akinesia. Indeed, we observed reduced movement in open field environment in PD mice, consistent with the theory of beta exaggeration in promoting akinesia.

In addition to persistent beta oscillation, transient fluctuations of beta are known to be critical for moment-to-moment regulation of movement^67,99,101^. Particularly relevant to gait regulation, Louie et al recently showed that beta and theta dynamics are temporally modulated by the gait cycle, and different human subjects exhibit distinct frequency signatures within the theta-beta frequency range of 5-15 Hz^26^. While our studies are not sufficiently powered to examine such changes, future studies by recording over an extended period of time would provide insights on how DA loss impacts the fluctuation of beta oscillations during gait cycles. Even though it is unclear how DBS selectively reduces individual STN neuron beta rhythmic spiking, the general reduction in firing rate during DBS suggests a reduced neuronal excitability using our DBS protocols, e.g. by reducing excitatory inputs or increasing membrane conductance (shunting inhibition), which could interfere with STN response to beta-rhythmic inputs. Indeed, using cellular membrane voltage imaging, we recently demonstrated that DBS alters membrane potential and interferes with cellular responding to inputs^73^. It is conceivable that DBS could alter STN neuron’s membrane potential, thereby reducing their responses to pathologically exaggerated beta-rhythmic inputs from GPe or other cortical and subcortical areas. While extracellular electrodes cannot capture synaptic inputs, future cellular voltage imaging studies could probe DBS mediated membrane biophysical effects underlying the observed beta-rhythmic spiking changes.

### DA-loss increases STN population gamma rhythmicity during movement, without altering individual neuron gamma-rhythmic firing

We found that positively modulated STN neurons exhibit a shift from beta-rhythmic to gamma-rhythmic firing during movement in both healthy and PD mice. Increased gamma synchrony between STN and cortex has been shown to be important for motor preparation during hand gripping^46^. The observed shift in individual neuron gamma dynamics is in line with the general function of STN gamma in promoting movement. As PD mice show an abundance of positively movement modulated neurons, the increase in STN synchrony during movement is also consistent with augmented firing gamma frequencies during movement.

While we detected a significantly greater gamma rhythmicity across the STN neuron population in PD mice, individual neuron’s gamma-rhythmic firing pattern remained constant between healthy and PD mice. As there is a higher proportion of positively movement modulated neurons in PD mice, the observed elevation of STN population gamma may be explained by the exaggerated activation of individual neurons rather than a shift in firing dynamics at the single cell level. Increased cortical gamma dynamics, and increased gamma synchrony between the STN and the cortex have been reported in PD^46,102–104^. However, previous LFP studies cannot distinguish the mechanism of such gamma enhancement. By recording many STN neurons simultaneously, we demonstrated that this increase in population STN gamma after DA loss is related to the exaggerated activation of STN neurons during movement, but not changes in individual neuron firing patterns. Interestingly, even though DBS reduced beta-rhythmic firing at rest in PD mice, it did not alter STN gamma. It is possible that DBS mediated cellular changes are not sufficient to bias gamma firing when STN neurons are in a more excited state with a high firing rate during movement.

### In PD mice, synchrony was increased during movement, and DBS reduced synchrony at rest

Synchrony in the STN has traditionally been measured by local LFP, which is thought to be dominated by synaptic activity rather than spiking^90^. Here we present spike synchrony between simultaneously recorded STN neuron pairs. While the mechanism leading to the emergence of sustained synchronization is complex, synchrony between STN neurons and motor related cortical beta oscillations play an important role in disrupting normal movement in PD ^105,106^. One possible explanation is that DA loss may reduce the strength of inhibitory inputs from GPe, causing the STN to be in a more excitable state and therefore respond more strongly to excitatory cortical inputs. The mechanism by which DBS reduces synchrony is again complex and poorly understood. DBS may suppress the hyperdirect pathway, shifting synaptic balance towards inhibition and causing information lesion^89,72^. Neuronal activity is theorized to be excited or suppressed by DBS depending on brain regions and synaptic inputs^107^. In the STN, high frequency DBS has been shown in brain slices to exaggerate inhibitory inputs^91^, which could reduce synchrony within the STN. This would explain why synchrony was comparable between healthy and PD mice at rest, but during movement the level of synchrony was elevated. While we cannot record the real time effect during DBS due to electrical stimulation artifact, our results demonstrate changes in spike synchrony in response to DA loss, and during intermittent DBS that is consistent with behavioral improvement.

## Methods and Materials

### Animal Preparation

All animal experiments were performed in accordance with the National Institute of Health Guide for Laboratory Animals and approved by the Boston University Institutional Animal Care and Use and Biosafety Committees. Same-sex mice from the same litters were generally housed together prior to surgery and single-housed post-surgery. Enrichment in the form of Igloos and running wheels was provided. Animal facilities were maintained around 70 °F and 50% humidity and were kept on a 12 h light/dark cycle. Mice were housed in reverse light cycle facilities to allow for recordings during their dark cycle, when they are more active.

The study preliminarily included 20 adult C57BL/6 mice, aged 8-16 weeks at experiment onset (Jackson Laboratory #000664, 12 females, 8 males). Data analysis was performed on 12 mice (8 female, 4 male), while 6 mice were excluded based on a histological inability to verify the STN, and 2 were additionally excluded due to lack of movement during baseline or DBS recordings, and therefore were excluded from further analysis.

To allow for head-fixed recording, mice were first surgical implanted with a headbar and marked with a craniotomy for STN target. Surgical procedures were largely identical to that described previously^108–110^. Briefly, mice were anesthetized with 1–3% isoflurane, and a 2 mm craniotomy was made over the STN (AP: -1.98, ML: 1.55, DV: -4.5). Surgical silicone adhesive (Kwik-Sil, World Precision Instruments) was placed on the craniotomy until the time of STN recordings to maintain hydration on the brain surface. Exposed skull regions were reinforced with Metabond Quick Adhesive Cement (Parkell, S380), a ground pin was placed in the cerebellum, and a custom aluminum headbar was affixed with dental cement (Stoelting, 5145). All mice received 72 hours of postoperative analgesia via a single preoperative intramuscular injection of sustained-release buprenorphine (0.03 mg/kg; Reckitt Benckiser Healthcare).

During the same surgery, half of the mice also received 6-OHDA injection in the medial forebrain bundle 1 week prior to headplate and craniotomy surgery. Specifically, desipramine was intraperitoneally delivered at 25 mg/kg, 30 minutes prior to the intracerebral injection of 6-OHDA to minimize 6-OHDA off target effects. 6-OHDA hydrobromide (Sigma Aldrich), was prepped in PBS with 0.1% ascorbate at a concentration of 5mg/ml. During survival surgery, 0.2 µl of prepared 6-OHDA was injected into the medial forebrain bundle (AP: -1.2 mm, ML: +1.1 mm, DV: -5 mm), to result in 1 µg delivered per mouse. One sham mouse was injected with 0.02% ascorbate/saline solution and was behaviorally and histologically indistinguishable from healthy mice, and therefore was included in the healthy control group. Lactated Ringers was delivered intraperitoneally for post-operation care for the week following injection to avoid dehydration.

Upon recovery from surgery, animals were first habituated for at least three sessions, before the electrophysiological recordings. During habituation, mice were head-fixed on the disk treadmill for increasingly longer periods (5 min-1 hour per day). Electrophysiological recordings were then performed once the mice were comfortable walking on the disk.

### Open field locomotion testing

To analyze the locomotion of mice in an open field environment, we placed mice into a custom 12”x12” open field box with a transparent base (**Sup. Fig. 1D-E**). Mice were video recorded with a Hero10 video camera (GoPro) positioned directly under the box recording 1080 pixels and 60 fps. A deep learning network in DeepLabCut was trained to capture all four paws. From this, the percent of time moving could be calculated. The number of ipsilateral and contralateral rotations was counted manually.

### Limb tracking and gait analysis under the head fixed condition

To analyze gait, paw positions were video recorded with a Hero10 video camera (GoPro) positioned directly under the mouse and the transparent disk was illuminated from below. Video frames were collected at a resolution of 1080 pixels at 60 fps. A deep learning network in DeepLabCut^80^ was trained on clips of six mice until a high degree of accuracy was achieved. The output of the tested videos was manually curated to detect for poor detection and retrained as necessary to achieve 90% of frames having an accuracy above 80%. All four paws were tracked throughout all the videos. Multi-animal training was utilized to enable DeepLabCut to take into account the orientation of limbs to one another, resulting in higher levels of accuracy.

We first excluded frames where paw identification accuracy by DeepLabCut was below 90%. We then applied a median filter on times below threshold to approximate where the paw was located. Next, the signal was demeaned and converted to centimeter scale using a fixed conversion factor determined by the fixed position of the camera relative to the disk. Next, a bandpass filter with cutoff frequencies from 0.5-30 Hz was applied to the position trace to remove any drift or high frequency noise. To identify movement periods, a moving standard deviation was thresholded above 0.25 cm to determine when gait bouts started and ended. Any gait bouts within 2 seconds of one another were merged and considered a single bout. Times at rest were defined as any period without limb movement. Steps were defined using the findpeaks function to find peaks and troughs of limb position, and manually inspected for accuracy. Any bout containing less than three steps were additionally excluded from analysis as either movement or rest periods. All movement traces were inspected and curated manually. Step times defined as the peaks of the trace, step duration is defined by the time from each step to the subsequent step time. Stride length was defined as the height of the swing length (trough to peak). An LED (Wurth Elektronik, 151051RS11000) was positioned in frame of the video camera, and at the start of every recording an LED pulse triggered by an Arduino Uno sent a pulse to the OpenEphys board to be recorded for aligning neural data to mouse movement.

### Electrical recording with custom linear electrode arrays

Silicon microelectrodes, such as the probes used in this proposal, are optimized for recording single neurons^111–113^. A custom silicon probe was designed to record and stimulate the STN (Torpedo Therapeutics Inc). Each recording sites are 11 µm x 8 µm, and the probe contain 19 sites at a 20 µm pitch. Each probe additionally features six circular 37 µm diameter stimulating electrodes (10 µm separation) positioned at the center of the recording sites. The custom silicon probe additionally has 13 recording sites at ∼700 µm above the STN sites, which were not used in the study. Probe fabrication broadly followed a previously published procedure^114^ with two notable differences. First, rather than the original method of electron beam lithography, the electrode geometry and routing are patterned using standard UV lithography using a maskless aligner (MLA-150, Heidelberg, Germany). Second, a layer of titanium nitride (TiN) was deposited over the electrode surface via sputtering from a TiN target. The resulting TiN structure consists of nanorods with pyramid-shaped caps, which significantly increase the electrode surface area, leading to lower impedance and enhanced charge storage capacity. For these experiments, we developed custom silicon probes to simultaneously target the subthalamic nucleus (stimulation and recording) and thalamus (recording only). The probes feature single penetrating shanks with two linear sets of recording electrodes targeting the STN (proximal to the shank tip) and thalamus (distal to the tip) and six stimulating electrodes located near the tip of the probe. Only data from the STN was included for this study.

During each recording session, mice were head-fixed over the clear disk treadmill. To prevent the mouse from damaging the electrode, a tail guard was placed to prevent the mouse from disturbing the probe. We first exposed the previously created craniotomy, gently removed dura removed with forceps. To track the electrode location, we placed 2 µl of dye (CellTracker CM-Dil, ThermoFisher) on the surface of the brain, inside the craniotomy, and lowered the tip of the probe to stain it. While inserting to the depth of the STN, the dye marked the location of the electrode tract to allow for visualization in histology.

All electrophysiological recordings were recorded with OpenEphys^115^ GUI and acquisition board (Open Ephys, 3^rd^ Generation). Briefly, the probe was lowered, zeroed at the skull surface, and lowered slowly until approximately 4mm depth. The probe was then advanced at 10 µm/s to look for tonically active neurons firing at ∼10-30 Hz ^51,75–78,116^, following the thin Zona Incerta area. Once multiple tonically active neurons were found, the brain was allowed to settle for 10 minutes before electrophysiological recordings began.

### DBS and simultaneous recording

DBS pulses were delivered using an isolated pulse stimulator (A-M Systems, Model 2100), with the timing of each stimulation pulse controlled by an Arduino Uno. DBS was delivered at 140Hz for 3 s every 13 s, for a total of 3 minutes (3.4 ± 0.66 mins across 12 mice). Stimulation pulses was bipolar square waves, 30 µA, 100-350 µs pulse widths (202 ± 104 µs). The ground pin in the cerebellum was used for the reference ground for stimulating and recording. To minimize stimulation artifact on recording electrodes, we used the built-in blanking circuits of the OpenEphys board to momentarily remove the Intan RHD Amplifiers during each DBS pulse train from the recording sites. The exact timing of the blanking circuits was controlled by the same TTL pulses used to command stimulation pulses through Arduino Uno. The blanking duration was experimentally determined to reduce the electrical artifact caused by stimulating and recording simultaneously while being as short as possible to record spiking soon after each stimulation pulse (2.8 ± 0.47 ms).

DBS pulses were kept at 30 µA per site in all mice to keep electrical field constant and to minimize electrical stimulation artifacts on our recordings. To account for the variation between electrodes, electrode placement, and intrinsic tissue excitability across mice, we empirically determined the DBS pulse width at the start of each experiment to match strength of stimulation to behavioral response. Specifically, we first determined the threshold pulse width that would evoke an immediate muscle twitch by gradually increasing the pulse width from 10 µs with a step size of 10 µs. We then used 75% of the threshold pulse width for all DBS experiments, and confirmed a lack of obvious muscle twitches change during stimulation.

### Spike Sorting

While blanking the Intan RHD amplifiers during DBS minimized stimulation artifact on recordings, saturation of the amplifier can remain take up to 5 ms after stimulation offset. Thus, we first identified and removed the recording data immediately after DBS offset with absolute value greater than 5 mV (i.e. data that lies well outside the ± 5 mV rated linearity range of the amplifiers). We then parsed each recording into continuous sections of valid data (i.e., neither blanked nor saturated) that ranged in duration from approximately 7.1 ms (the period of 140 Hz stimulation) to several minutes (entire recordings where no stimulation was applied). We then applied a third order Savitzky-Golay filter to each section (5 ms time constant; implemented using the *savgol filter* function of the scipy package) and subtracted the result from the raw data. Similarly to the SALPA algorithm^117^, this method removes both low-frequency spectral content of the LFP and any remaining settling artifacts caused by amplifier saturation in a single processing step. We then apply common-average re-referencing to the data obtained from the proximal and distal groups of recording sites, respectively. Finally, we re-combine the processed data sections and linearly interpolate missing samples. In contrast to classical time-domain filtering^118^, this approach avoids signal contamination from any sharp transitions at the edges of the blanking period. We then re-referenced the data of each electrode site to the mean of all electrode sites. Finally, we re-combine the processed data sections and linearly interpolate missing samples.

We grouped and concatenated preprocessed recordings by electrode depth and used a lightly modified version of the Kilosort4 algorithm to identify putative single- and multi-units^119^. Specifically, the main algorithm was untouched, but common average re-referencing and high-pass filtering were disabled as they had already been applied at the preprocessing stage. Finally, spikes were manually curated using the Phy package^120^ and the quality of the analysis was confirmed using custom visualization tools (based on the *matplotlib*, *seaborn*, *pyqt*, *pyqtgraph* and *ephyviewer* packages).

### Movement-modulated neuron bootstrapping classification

To determine whether a neuron was modulated by movement, we compared the observed firing rate during previously identified movement periods to that of randomly shuffled ISIs 1000 times. If the observed proportion of spikes during movement was above the highest 95^th^ percentile of shuffled distributions, the neuron was classified as “positive modulated”. If the observed proportion was below the lowest 95^th^ percentile, the neuron was classified as “negative modulated”. If neither was true, the neuron was classified as “non-modulated”.

### STN neuron spatial distribution

The depth of each STN neuron recorded was approximated by the location of the recording site with the highest amplitude waveforms of a given neuron. The dorsal-ventral location within the STN was then approximated by the recorded depth of the electrode relative to the brain surface and the location of the site on the electrode.

### Inter-Spike Interval analysis

Inter-Spike-Interval (ISI) return maps were generated by taking all the ISIs from a neural recording, and plotting each spike coordinate as ISI(n) versus ISI(n-1). We then binned the generated data into a 100x100 grid spanning 1 ms to 1000 ms ISI values. To calculate population averages, the histogram was normalized by the highest proportion observed, and averaged across neurons.

To compare whether populations had a higher proportion of beta band or gamma frequency band firing, we calculated the proportion of each band for each neuron. This value of each proportion was calculated by finding the number of ISIs falling within the bands of 13-30 Hz and 30-100 Hz, and diving by the overall number of ISIs found per neuron. We then compared the proportions across groups (baseline versus DBS). This provided an estimate for how beta and gamma frequency firing was impacted.

### Gait modulation of neurons using limb-spike phase locking

The phase of gait traces was calculated by taking the angle of the Hilbert Transform of the trace, after being filtered through a bandpass filter (cutoff 0.5-4Hz). Spike times were only included during movement as defined previously, and the spike phase was then determined by the limb phase at the spike event.

### Cross-correlogram between two neurons

To create the correlation per lag probability correlograms, we took the difference in spike times between two neurons’ rasters, and plotted the histogram of differences (50 bins from -25 to 25ms lag). The values were normalized by the overall number of spikes per recording. For visualization, the histogram was smoothed by a moving-mean window size of 2 values.

We estimated STN network synchrony through computing pair-wise cross correlation between all pairs of neurons with >1 Hz spike rate. In order to calculate cross correlation between two spike rasters, we convolved them with 50 ms time windows. The average value of the cross correlation between all neuron pairs resulted in the average synchrony used for **Figure 6**. For DBS trials, so as to not be biased by the inability to spike sort during DBS, we omitted times of DBS when calculating cross correlation. As we know that DBS reduces firing rate significantly compared to baseline, we used a second metric of synchrony to validate our results. Synchrony calculations were repeated using Spike Time Tiling Coefficients^121^, a measure known to reduce the effects of variable spike rates, and resulted in the same trend as calculated by cross correlation (**Sup Fig. 6**).

To account for any changes in spike rates over time, STTC values were calculated between neurons pairs, by evaluating the equation below, using the parameters: T_A_: The proportion of total recording time which lies within ±Δt of a spike from neuron A. T_B_: calculated similarly from neuron B. P_A_: the proportion of spikes from A which lie within ±Δt of any spike from B. P_B_: Calculated similarly.

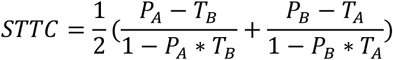

### 6-OHDA lesion quantification and histology

Animals were perfused at the completion of the study, their brains extracted, post fixed in 4% paraformaldehyde, and sliced into 50μm coronal slices. Electrode tract location confirmation was performed using the CellTracker Dye applied to the electrodes prior to implantation, and imaged using a fluorescent microscope (Olympus VS120 Virtual Slide Scanner). Mice without clear electrode tract through the STN were excluded from the study.

Dopamine loss was assessed by immunofluorescence of tyrosine hydroxylase (TH) on 6 slices spaced 100-200μm apart, spanning the striatum. Striatal brain slices were stained using a Rabbit anti-TH antibody (Millipore Sigma, No. AB152, 1:500) and a Mouse anti-Neuronal Nuclei antibody (Millipore Sigma, No. MAB377), followed by Alexa Fluor 568 goat anti-rabbit secondary antibody (Invitrogen, No. A11011, 1:1000), Alexa Fluor 488 goat anti-mouse secondary antibody (Invitrogen, No. A10680, 1:500), and Hoechst 33342 DNA-specific fluorescent stain (ThermoFisher, No. 62249, 1:1000). The stained slices were imaged using Olympus VS120 slide scanner microscope at 10× magnification to visualize Alexa Fluor 568, Alexa Fluor 488, and Hoechst fluorescence. Each image was then automatically tiled, and the lesioned side was confirmed using the Alexa Fluor 568 fluorescence. Striatal TH staining intensity was calculated for both the lesioned hemisphere (TH_lesion_) and the intact hemisphere (TH_healthy_). The percent reduction of dopamine innervation was defined as:

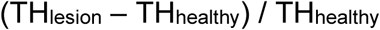

### Statistics

To test whether number of rotations, stride lengths, or stride durations changed between individual mice or individual recordings, we used non parametric Kruskal-Wallis Test to account for non-normal distributions of movement parameters. Post hoc pairwise comparisons were performed using Dunn’s multiple-comparison procedure with Bonferroni correction.

To test whether stride lengths of healthy and PD populations across baseline and DBS trials showed differences, we estimated a pseudo-Poisson generalized linear mixed-effects (GLME) model using the following equation:

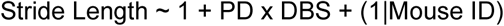

where disease status (PD vs Healthy), DBS condition (on vs off), and their interaction were included as fixed effects, and with mouse identity included as a random intercept to model between-animal variability. This model was repeated for stride duration. We estimated the difference in stride limb asymmetry in individuals and populations using the Mann-Whitney U test between adjacent limbs of mice.

To assess the difference in firing rate between healthy and PD mice, and at rest versus during movement, we used a non-parametric Kruskal-Wallis test, to account for non-normal distributions of firing rate. Post hoc pairwise comparisons were performed using Dunn’s multiple-comparison procedure with Bonferroni correction. Further post hoc comparison used Wilcoxon Signed-Rank to compare the same neurons across movement and rest, as the data were paired. For estimating the difference between counts of movement modulated neurons across PD and healthy mice, we used Chi-Square tests.

To test the changes of firing patterns using populations of ISIs, we compared healthy and PD median ISIs, using the Kolmogorov–Smirnov (KS) test to test whether two cumulative distribution functions are significantly different. When comparing median ISI distributions between movement and rest, Wilcoxon Signed-Rank test was used for paired data. To measure the proportion of ISIs in a frequency band between groups (beta 33 ms-76 ms or gamma 10-33 ms), we recorded the proportion of ISIs that fell within a frequency range for each neuron, and compared across populations using the Mann-Whitney U test. The same test was performed on distributions between healthy and DBS trials.

To perform statistical analysis of spike-limb phase directionality, we used the Rayleigh Statistic to measure the directionality and power of spiking aligned to the phase of limb position. Across the population of neurons, we then compared the average power and direction of phase using the non-parametric Watson Williams test. Fisher’s exact test was used to compare any two counts of gait modulated categories. All polar statistics were calculated using the Circular Statistics Toolbox (Directional Statistics) Version 1.21.0.0 for MATLAB.

To assess the levels of synchrony across DBS and baseline recordings, we utilized the Wilcoxon Signed-Rank test to compare the paired average Pearson’s Correlation values from all neurons in each recording, both for healthy and PD mice. This test was also employed for STTC comparisons. This test was consistently used for rest, motion, and whole recording synchrony. We then sought to compare the baseline synchrony of healthy and PD mice. In order to take into account mouse level variability in neuron count, we employed a pseudo-Poisson generalized linear mixed-effects (GLME) model using the following equation:

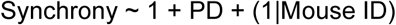

where disease status was (PD vs Healthy) was included as the fixed effect, and mouse identity as a random intercept to model between-animal variability. This model was chosen as it does not assume normality of observed data, and factors in variability between animals to assess inter-group synchrony changes.

## Acknowledgments

We thank members of Han Lab for their help throughout the study. X. H. acknowledges funding from the NIH (1RF1NS129520, 1R01NS119483, 1R01NS115797, and 1R01NS139524). I. v. W. and R. D. acknowledge NIH 5R44MH118155. The authors additionally acknowledge support from the Shared Computing Cluster in Boston University’s Research Computing Services, and Boston University Micro and Nano Imaging Facility (support by NIH S10OD024993).

## Contributions

L.S. Performed all electrophysiology experiments and analyzed the data. L.S. and A.L. prepared the animals for experiments and performed histology. H.T. provided technical assistance. R.D. assisted with data analysis and provided technical assistance. I. v. W. supervised all work at Torpedo Therapeutics Inc. that designed and fabricated probes, as well as provided technical assistance to all animal experiments. X.H. supervised all work at Boston University, including data collection and analysis. L.S. and X.H. wrote the manuscript. All authors edited the manuscript.

## Supplemental Figures

**Supplemental Figure 1:**
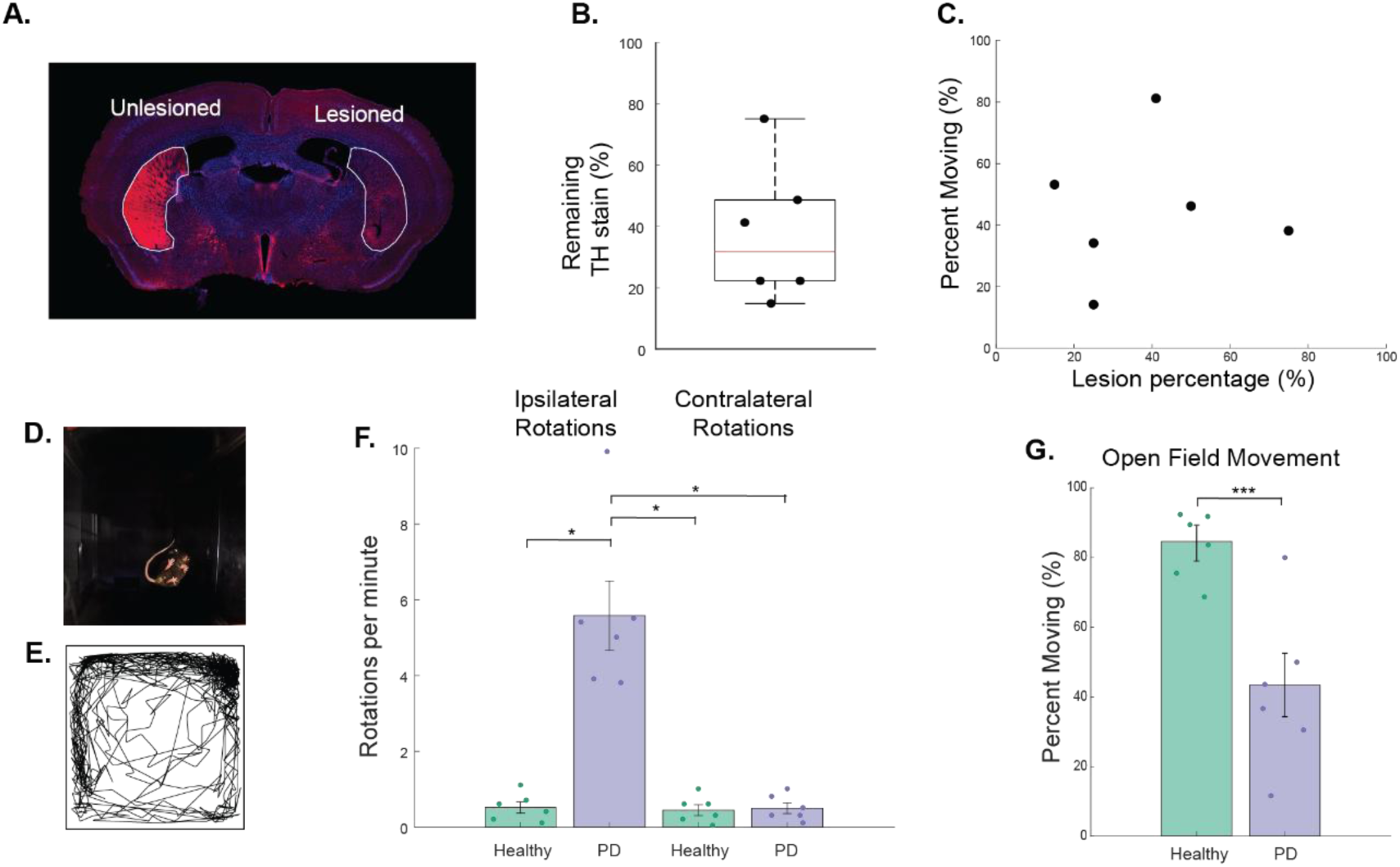
Histology and open field analysis. (**A**) An example coronal striatal slice showing DA loss. Blue: DAPI nuclear stain; Red: immunofluorescence against tyrosine hydroxylase (TH staining). Outlines highlight the striatum area used to quantify TH staining. **(B)** Quantification of DA loss, calculated as TH immunofluorescence of the lesioned hemisphere over the unlesioned hemisphere. Each dot represents a mouse. (**C**) DA loss versus the fraction of time moving. (**D**) An example image frame showing the four paws and body position of a mouse during the open field testing. (**E**) Position of the mouse in the open field environment as in D, during an example session. (**F**) Ipsilateral and contralateral rotations per minute for healthy and PD mice. (**G**) Percent of time moving in the open field in healthy and PD mice. Shown are mean ± standard deviation. Each dot is a mouse. *, p<0.05. **, p<0.01. ***, p<0.005, Wilcoxon rank-sum test.

**Supplemental Figure 2:**
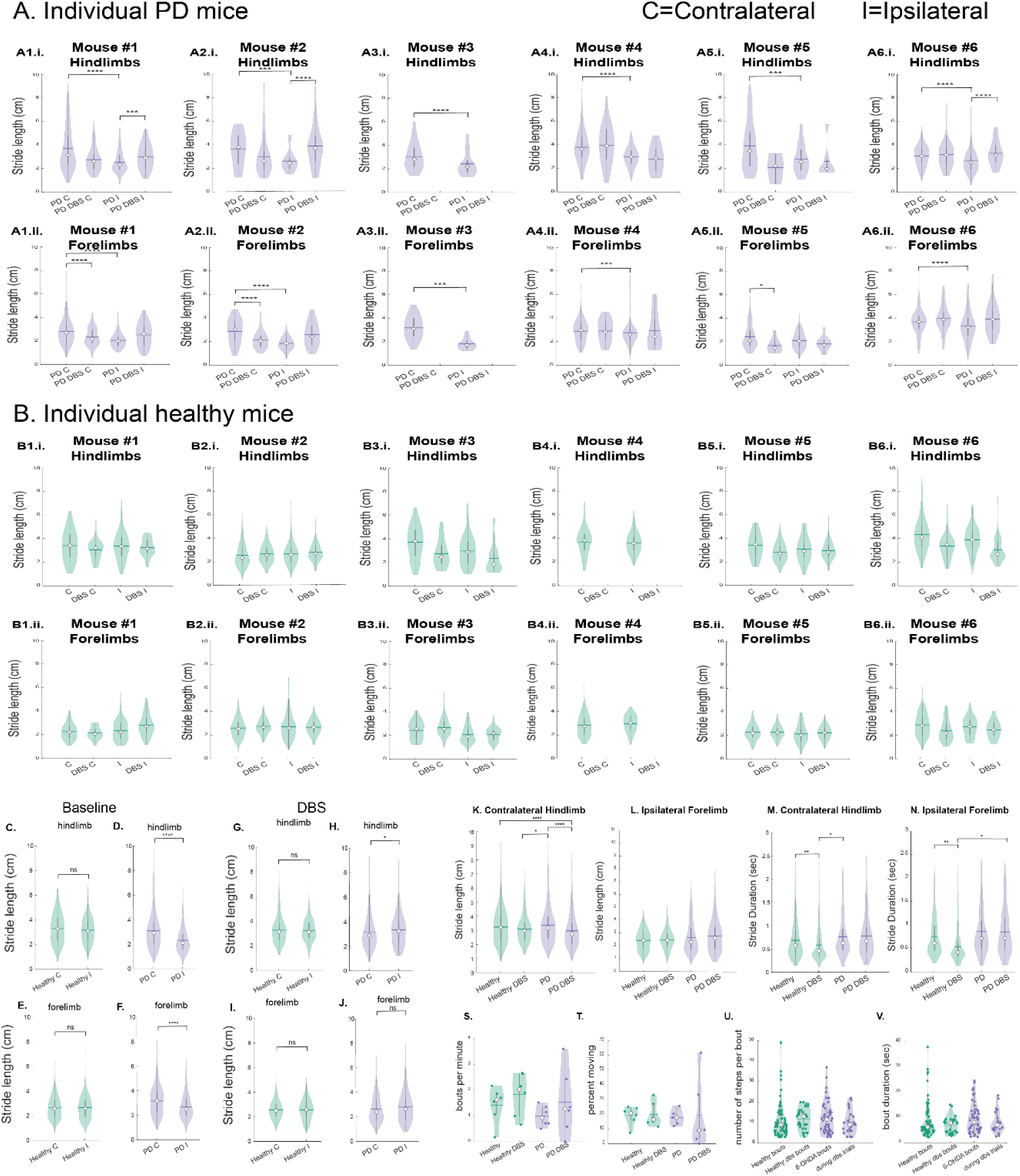
Comparison of limb parameters. (**A**) Comparison of stride lengths between the 2 hindlimbs (**Ai**) and 2 forelimbs (**Aii**) in each of the 6 PD mice (**A1-6**), during baseline without DBS (left) and during DBS (right). Mouse A3 had little movement during DBS and was excluded from the analysis of DBS effects. (**B**) Similar to A, but in each of the 6 healthy mice (**B1-6**). (**C, D**) Comparison of stride length of contralateral versus ipsilateral hindlimbs in healthy (**C**) and PD (**D**) population during baseline period. (**E, F**) Similar to **C** and **D**, but for forelimbs. (**G-J**) Similar to **C-F**, but during DBS periods. (**K**). Comparison of the stride length of contralateral hindlimb between PD and healthy population, and with versus without DBS. (**L**) Same as **K**, but for ipsilateral forelimb (**L**). (**M, N**), Similar to **K** and **L**, but for stride durations. (**S**) Comparison of bouts per minute between healthy and PD population, during baseline versus DBS. (**T-V**) Similar to **S**, but for percent of time moving. (**T**), number of steps per bout (**U**) and bout duration (**V**). *, p<0.05. **, p<0.01. ***, p<0.005. ****, p<0.001. Wilcoxon rank-sum for **A-B**, **K-V**. Wilcoxon signed-rank for **C-J**.

**Supplemental Figure 3:**
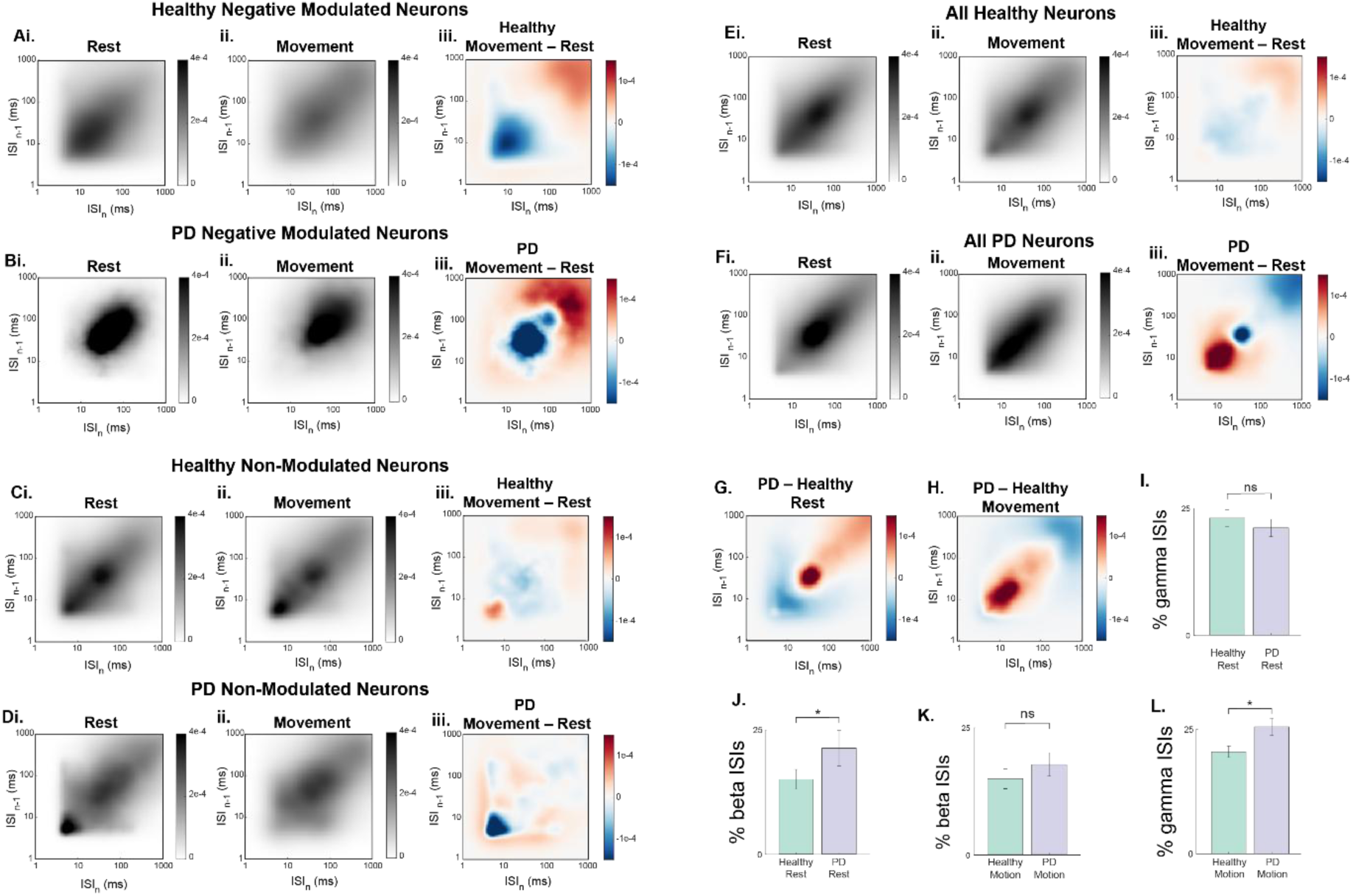
ISI distributions of neurons in healthy and PD mice. (**A**) ISI return map of negatively modulated neurons in healthy mice at rest (**Ai**), during movement (**Aii**), and their difference (**Aiii**, shown as **Aii** minus **Ai**). N=22 neurons. (**B**) Similar to A, but for negatively modulated neurons in PD mice. N=5 neurons. (**C, D**) Similar to A and B, but for non-modulated neurons in healthy mice (**C**) and PD mice (**D**). N=35 healhty and N=9 PD neurons. (**E, F**) Similar to A and B, but for all neurons regardless of movement modulation in healthy mice (**E**) and PD mice (**F**). (**G, H**) Difference in the ISI return map between healthy and PD mice (PD minus healthy) at rest (**G**) and during movement (**H**). (**I, L**) Percentage of gamma ISIs in ISI distributions, across healthy and PD groups, during rest (**I**) and movement (**L**). (**J, K**) Percentage of beta ISIs in ISI distributions, across healthy and PD groups, during rest (**J**) and movement (**K**). *, p<0.05, Wilcoxon rank-sum.

**Supplemental Figure 4:**
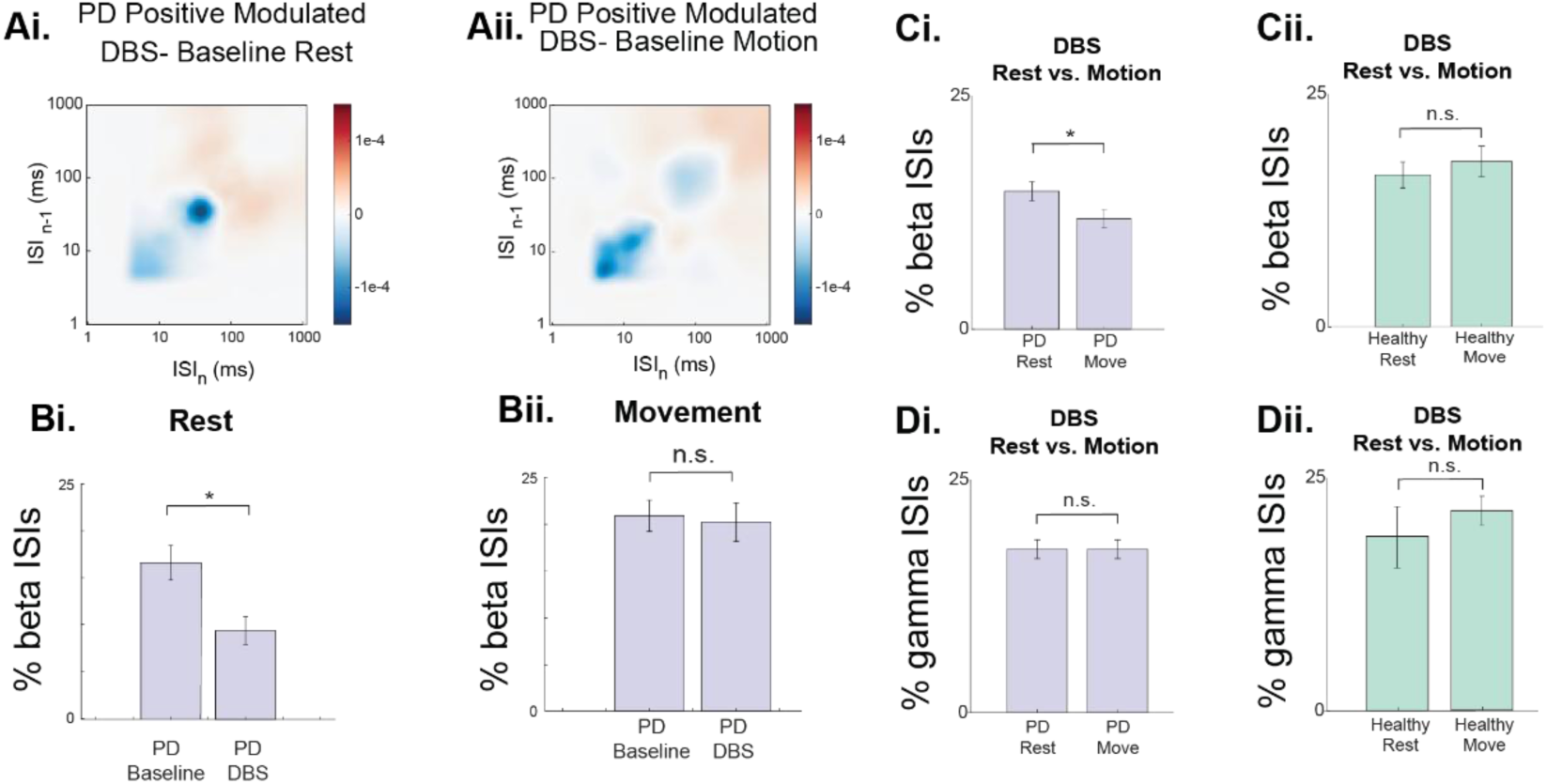
DBS reduces beta-rhythmic firing in PD mice. (**A**) Difference in the ISI return map of positively modulated neurons between baseline and DBS (DBS-baseline) in PD mice at rest (**Ai**) and during movement (**Aii**). **(B)** Percent of beta ISIs between baseline and DBS in PD mice at rest (**Bi**) and during movement (**Bii**). (**C**) Comparison of the fraction of beta ISIs between rest and movement in PD mice (**i**) and healthy mice (**ii**). **(D)** Same as **C**, but for gamma ISIs. *, p<0.05, Wilcoxon signed-rank.

**Supplemental Figure 5:**
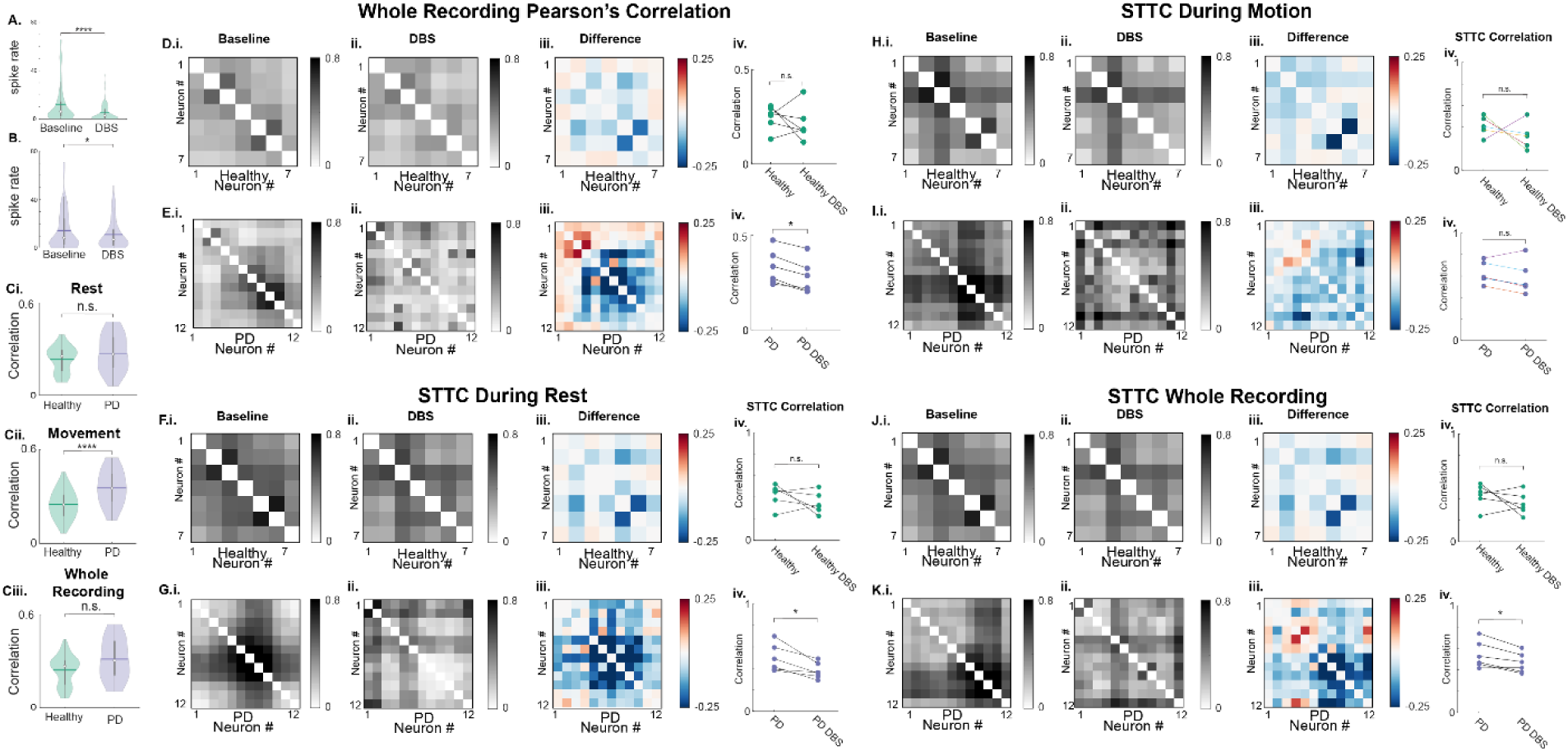
Spike synchrony computed by Pearson’s Correlation and Spike Time Tiling Coefficient (STTC). (**A, B**) Spike rate across all neurons during the entire recording under baseline and DBS conditions in healthy mice (**A**) and PD mice (**B**). (**C**) Correlation strength across neurons in PD and healthy mice (n=88 neurons in healthy, n=76 in PD, GLME ****, p<0.001). Correlation strength of each neuron is computed by the mean of the correlation coefficients between the given neuron and all other simultaneously recorded neurons during rest (**i**), during movement (**ii**), and during the entire recording (**iii**). (**D**) Correlation coefficients of all neuron pairs from an example recording session of a healthy mouse during baseline period (**i**), during DBS (**ii**), and their difference (DBS-Baseline). (**Div**) Averaged correlation coefficients of individual neurons in healthy mice during baseline period and DBS. Each data point represents the average value of all neuron pairs in a recording. (**E**) Similar to **D**, but for a recording session in a PD mouse. (**F**) Similar to **D**, **E**, but with spike synchrony during the whole recording session computed by STTC instead of Pearson’s Correlation. (**H**, **I**), similar to **F**, **G**, but during rest. (**J**, **K**), similar to **F**, **G**, but during movement. *, p<0.05, **, p<0.01. ***, p<0.005. ****, p<0.001, Wilcoxon signed-rank

